# Direct *in vivo* mapping of functional suppressors in glioblastoma genome

**DOI:** 10.1101/153460

**Authors:** Ryan D. Chow, Christopher D. Guzman, Guangchuan Wang, Florian Schmidt, Mark W. Youngblood, Lupeng Ye, Youssef Errami, Matthew B. Dong, Michael A. Martinez, Sensen Zhang, Paul Renauer, Kaya Bilguvar, Murat Gunel, Phillip A. Sharp, Feng Zhang, Randall J. Platt, Sidi Chen

**Author notes:** Co-first authors (Ryan D. Chow, Christopher D. Guzman, Guangchuan Wang, Florian Schmidt). Co-second authors (Mark W. Youngblood, Lupeng Ye). Correspondence: RJP, SC, +1*-203-737-3825 (office)*, +*1-203-737-4952 (lab).

## Abstract

Glioblastoma (GBM) is one of the deadliest cancers, with limited effective treatments and single-digit five-year survival ^1-7^. A causative understanding of genetic factors that regulate GBM formation is of central importance ^8-19^. However, a global, quantitative and functional understanding of gliomagenesis in the native brain environment has been lacking due to multiple challenges. Here, we developed an adeno-associated virus (AAV) mediated autochthonous CRISPR screen and directly mapped functional suppressors in the GBM genome. Stereotaxic delivery of an AAV library targeting significantly mutated genes into fully immunocompetent conditional Cas9 mice robustly led to gliomagenesis, resulting in tumors that recapitulate features of human GBM. Targeted capture sequencing revealed deep mutational profiles with diverse patterns across mice, uncovering *in vivo* roles of previously uncharacterized factors in GBM such as immune regulator *B2m,* zinc finger protein *Zc3h13,* transcription repressor *Cic,* epigenetic regulators *Mll2/3* and *Arid1b,* alongside canonical tumor suppressors *Nf1* and *Pten*. Comparative cancer genomics showed that the mutation frequencies across all genes tested in mice significantly correlate with those in human from two independent patient cohorts. Co-mutation analysis identified frequently co-occurring driver combinations, which were validated using AAV minipools, such as *Mll2, B2m-Nf1*, *Mll3-Nf1* and *Zc3h13-Rb1*. Distinct from *Nf1*-oncotype tumors, *Rb1*-oncotype tumors exhibit undifferentiated histopathology phenotype and aberrant activation of developmental reprogramming signatures such as *Homeobox* gene clusters. The secondary addition of *Zc3h13* or *Pten* mutations drastically altered the gene expression profiles of *Rb1* mutants and rendered them more resistant to the GBM chemotherapeutic temozolomide. Our study provides a systematic functional landscape of GBM suppressors directly *in vivo*, opening new paths for high-throughput molecular mapping and cancer phenotyping.

## Introduction

Glioblastoma (Glioblastoma multiforme, GBM) is one of the deadliest cancers, with a single-digit five-year survival rate ^1^. Current standard of care fails to cure the vast majority of patients with this disease ^1-3^, leaving them a median survival of 12.2 to 18.2 months, and a five-year survival of less than 10% ^4-7^. Genetic alterations are the major causes for the transformation of normal cells in the brain into highly malignant glioma cells ^1,8-10^, particularly those occurring in general or brain-specific oncogenes and tumor suppressor genes ^11-15^. The first genome atlas of GBM uncovered 453 validated non-silent somatic mutations in 223 unique genes, which were further refined to a total of 71 significantly mutated genes (SMGs) ^16^. Subsequent integrative genomic analyses revealed comprehensive mutational landscapes in GBM, uncovering 21 to 75 SMGs across multiple different cohorts of patients ^13,15-19^. Many of the newly discovered genes have never been characterized in GBM; thus, their functional roles in gliomagenesis remain largely unknown ^13,17^. Further complicating the interpretation of causality, mutations can occur in novel combinations across individual patients, leading to drastically different pathological features, prognoses, and therapeutic responses ^3,20-22^. Thus, a deeper functional understanding of gliomagenesis and a quantitative measurement of phenotypic effects across various combinations of drivers are both of central importance.

To date, no study has comprehensively and combinatorially investigated which of the mutations identified in human patients can indeed functionally drive GBM from normal cells in the brain, because achieving such a global, quantitative and functional understanding of gliomagenesis in a controlled experimental setting has been challenging ^17^. The major barriers include accurate delivery, precise genome manipulation, efficient massively parallel perturbation, and unbiased, high-sensitivity quantitative readout, all of which have to be achieved simultaneously in the native brain microenvironment. We overcame these challenges by developing and performing an AAV-mediated direct *in vivo* autochthonous CRISPR screen in the brain of fully immunocompetent mice, coupled with capture sequencing to achieve an ultra-deep readout of all functional variants. With this quantitative data, we identified multiple new drivers and co-occurring drivers, and subsequently validated a set of such combinations. Transcriptome profiling of these validated mutants revealed distinct expression signatures, either between genotypes or in response to temozolomide (TMZ) treatment. Utilizing this direct *in vivo* autochthonous screen approach, we mapped the functional landscape of GBM suppressors in the native microenvironment of the mouse brain.

## Results

To directly test the function of putative SMGs in the mouse brain, we set out to develop a direct *in vivo* autochthonous screening strategy, which necessitates pooled mutagenesis of normal cells directly in the native organ and subsequent deconvolution of mutant phenotypes. Because GBM is a disease originating from astrocytes, we generated an AAV-CRISPR vector that encodes Cre recombinase under a *Glial fibrillary acidic protein* (*GFAP*) promoter, resulting in conditional expression of Cas9 and GFP in astrocytes when injected into a conditional LSL-Cas9 mouse (Methods) (Figure S1A). The vector also contains an sgRNA targeting *Trp53*, with the initial intent to generate co-mutational *Trp53* knockout that might exhibit genome instability and thus be sensitized to tumorigenesis ^23-27^. Local viral delivery into the brain restricts the number of transducible cells, and cancer genomes generally consist of dozens to hundreds of SMGs ^28-31^. With these considerations in mind, we designed an sgRNA library (mTSG library) targeting the mouse homologs of top-ranked pan-cancer SMGs (Methods), plus 7 genes with essential molecular functions that we initially considered as internal controls (Figure 1A) (Table S1). We synthesized all sgRNAs as a pooled library, cloned them into the AAV-CRISPR vector at greater than 100x coverage, and deep-sequenced the library to ensure all sgRNAs were fully covered and represented with a tight lognormal distribution (99% within two orders of magnitude) (Figure 1A, Figure S1B). We generated high-titer AAVs (> 1 * 10^12^ viral particles per mL) from the plasmid that contained the mTSG library (AAV-mTSG), as well as the empty vector (AAV-vector) (Figure 1A). We then stereotaxically injected AAV-mTSG, AAV-vector or PBS into the lateral ventricle (LV, n = 40 mice) or hippocampus (HPF, n = 16 mice) in the brains of LSL-Cas9 mice (Methods). We performed magnetic resonance imaging (MRI) to scan the brains of these mice at four-months post-injection, and found that half (9/18 = 50%) of AAV-mTSG library transduced animals developed brain tumors at this time point, whereas none of the AAV-vector or PBS injected animals had detectable tumors by MRI (Figure 1B) (Figure S1E) (Table S2). Quantification of tumor volumes showed that AAV-mTSG transduced mice had average tumor volumes of 70.2 mm^3^ (including animals without tumors), or 140.3 mm^3^ (excluding animals without a tumor) (two-tailed Welch’s t-test, *p* = 0.018, mTSG vs. vector or PBS) (Figure 1C) (Table S2). These data suggested that the AAV-mTSG viral library robustly initiated and driven tumorigenesis in the brains of LSL-Cas9 mice.

**Figure 1.**
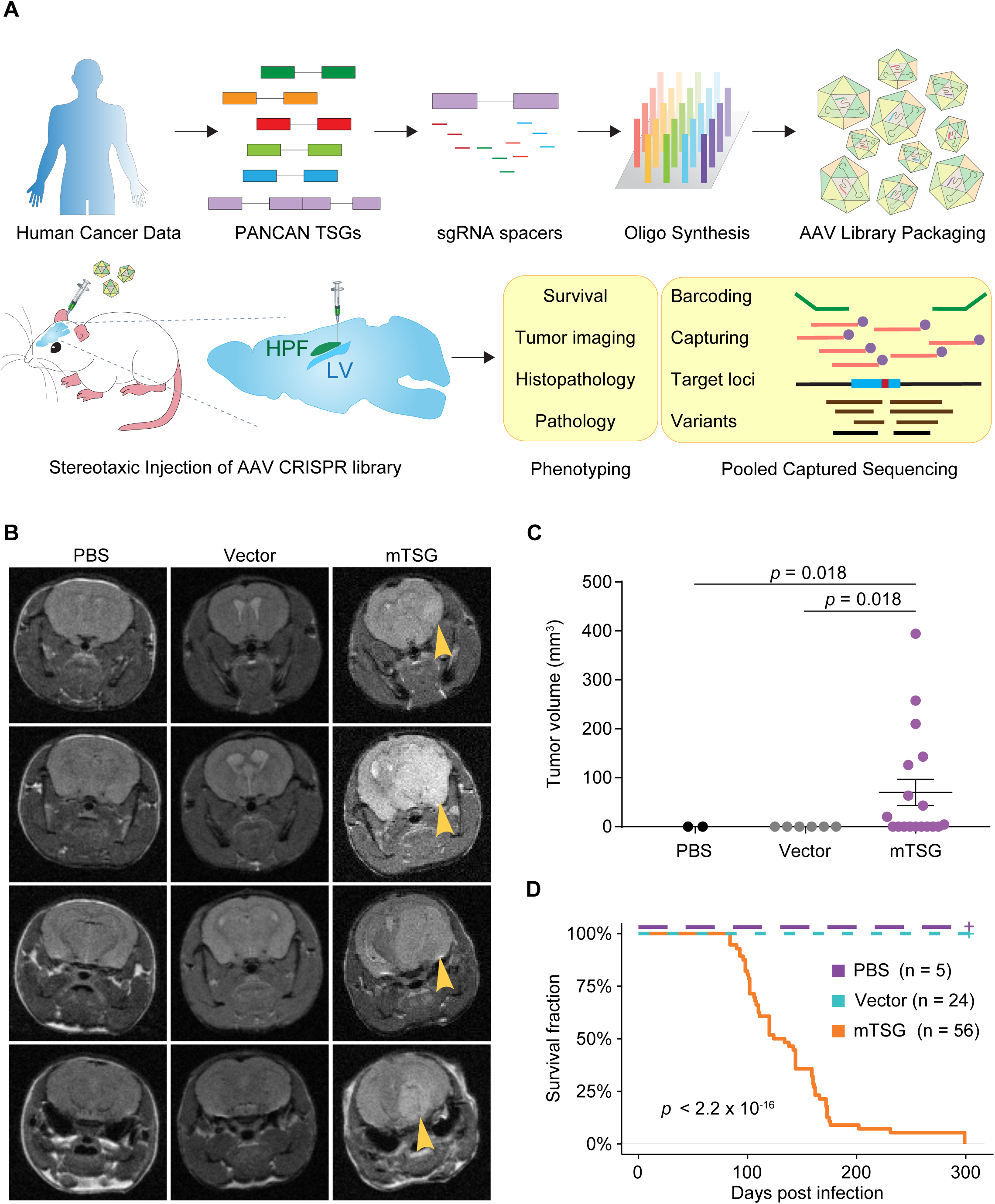
AAV-CRISPR library mediated direct mutagenesis induced potent autochthonous tumorigenesis in the mouse brain. **A.** Schematics of direct *in vivo* AAV-CRISPR GBM screen design. Top panel, AAV-mTSG library design, synthesis and production. A ranked list of TSGs was generated based on significantly mutated genes from TCGA pan-cancer analysis and the mouse orthologs were retrieved (mTSGs); sgRNA spacers were computationally identified, synthesized as a pool, and cloned into an AAV vector. Bottom panel, stereotaxic injection of AAV library and subsequent analysis. HPF, hippocampus; LV, lateral ventricle. SgRNAs (blue box) generate indel variants at genomic loci around predicted cutting sites (small red box), which were readout in pooled directly using barcoded captured high-throughput sequencing. **B.** MRI imaging of brains in the mice injected with PBS, AAV-vector or AAV-mTSG library. MRI sections showing brain tumors in AAV-mTSG injected mice, but not in and matching sections from PBS or AAV-vector injected mice. Arrowheads indicate brain tumors. **C.** MRI-based volumetric quantification of tumor size. Two-tailed t-test, *p* = 0.018, mTSG vs. vector or PBS; two-tailed t-test, *p* = 0.5, vector vs. PBS (PBS, n = 2; Vector, n = 6; mTSG, n = 18). **D.** Kaplan-Meier curves for overall survival (OS) of mice injected with PBS, AAV-vector or AAV-mTSG library. OS for PBS and vector groups are both 100%, where the curves are dashed and slightly offset for visibility. Log-rank (LR) test, *p* < 2.20 * 10^−16^, mTSG vs. vector or PBS; LR test, *p* = 1, vector vs. PBS.

We analyzed the overall survival of a cohort of LSL-Cas9 mice injected with AAV-mTSG, AAV-vector or PBS (Table S3). In this screen, injection location did not affect the rate of tumor development as reflected by overall survival (two sided Mann-Whitney U test of HPF vs LV, *p* = 0.054) (Table S3), and thus were considered as one group (AAV-mTSG). For the AAV-mTSG transduced group, the first three animals died 84 days post injection (dpi), 90% of animals did not survive 176 dpi, and all 56 AAV-mTSG transduced animals reached their survival endpoints within 299 days (*i.e.* died, or had a poor body condition score (BCS, < 2) thus were euthanized) (Table S3) (Figure 1D). The median survival time of the AAV-mTSG group was 129 days (95% confidence interval (CI) = 111 to 159 days) (Figure 1D), consistent with the presence of large tumors in half of the mice at 4 months by MRI. In sharp contrast, all 24 AAV-vector and all 5 PBS injected animals survived the whole duration of the study and maintained good body condition (BCS = 5) (log-rank (LR) test, *p* < 2.2 * 10^−16^, mTSG vs. vector or PBS; LR test, *p* = 1, vector vs PBS) (Figure 1D). For the vast majority (96.4%, or 54/56) of AAV-mTSG injected mice, macrocephaly was observed at the survival endpoint (Figure S1C), suggesting that they had developed brain tumors. On the contrary, macrocephaly was observed in none of the AAV-vector (0/24) or PBS (0/5) injected mice during the whole study (two-tailed Fischer’s exact test, *p* < 1 * 10^5^, mTSG vs. vector or PBS; *p* = 1, vector vs. PBS). These data indicated that the brain tumors induced by the AAV-mTSG viral library were typically lethal.

Under a fluorescent stereoscope, we observed that AAV-mTSG mice had sizable GFP-positive masses that deformed the brains (100%, or 6/6) (Figure S1D, Figure 2A). AAV-vector mice had diffuse GFP-positive regions in the brain with fully normal morphology, suggesting these were AAV-transduced cells expressing Cas9-GFP induced by Cre expression, which had not become tumors (n = 2) (Figure S1D, Figure 2A). PBS injected or uninjected mice had no detectable GFP expression even at long exposure (n = 3) (Figure S1D). Immunohistochemistry (IHC) analysis showed that AAV-mTSG induced tumors stained positive for Cas9 and GFP, consistent with them having arisen from cells with activation of Cas9-GFP expression (Figure 2A). These tumors were also positive for GFAP, an astrocytic marker (Figure 2A); and for Ki67, a proliferation marker (Figure 2A). AAV-vector transduced brains stained positive for Cas9 and GFP in a subset of cells at the injection site (Figure 2A), but these cells were not proliferative (Ki67 negative) and did not have tumor-like pathological features (Figure 2A). PBS injected mice stained negative for Cas9, GFP and Ki67 (Figure 2A). Endpoint histopathology showed that the vast majority of AAV-mTSG mice developed brain tumors (10/11 = 91%), whereas none of AAV-vector (0/7 = 0%) or PBS (0/3 = 0%) mice had detectable tumors (two-tailed Fischer’s exact test: *p* = 0.0003, mTSG vs. vector; Fisher’s exact test, *p* = 0.011, mTSG vs. PBS) (Figure 2A-B) (Table S4). The mean endpoint tumor size as measured by area in the brain sections for the AAV-mTSG group was 13.9 mm^2^, as compared to 0 mm^2^ in the two control groups (two-tailed Welch’s t-test, *p* = 0.0026, mTSG vs. vector or PBS) (Figure 2B). The brain tumors in AAV-mTSG mice showed pathological features of dense cellular structure with proliferative spindles, nuclear aneuploidy and pleiomorphism, giant cells, regions of necrosis, angiogenesis and hemorrhage (Figure 2C), all of which are hallmark features of human GBM ^2^. Clinical features such as deformation of the brain, invasion, loss of neuronal bundles, necrosis and hemorrhage were further corroborated by special staining methods such as Luxol fast blue Cresyl violet (LFB/CV), Wright Giemsa, Masson and Alcian blue Periodic acid – Schiff (AB/PAS) (Figure S2). We investigated a panel of human GBM clinical samples from Yale Glioma tissue bank, and confirmed the observation of these pathological features (Figure S3A-B). These data suggested that the AAV-mTSG library-induced autochthonous brain tumors recapitulate various histological and pathological features of human GBM.

**Figure 2.**
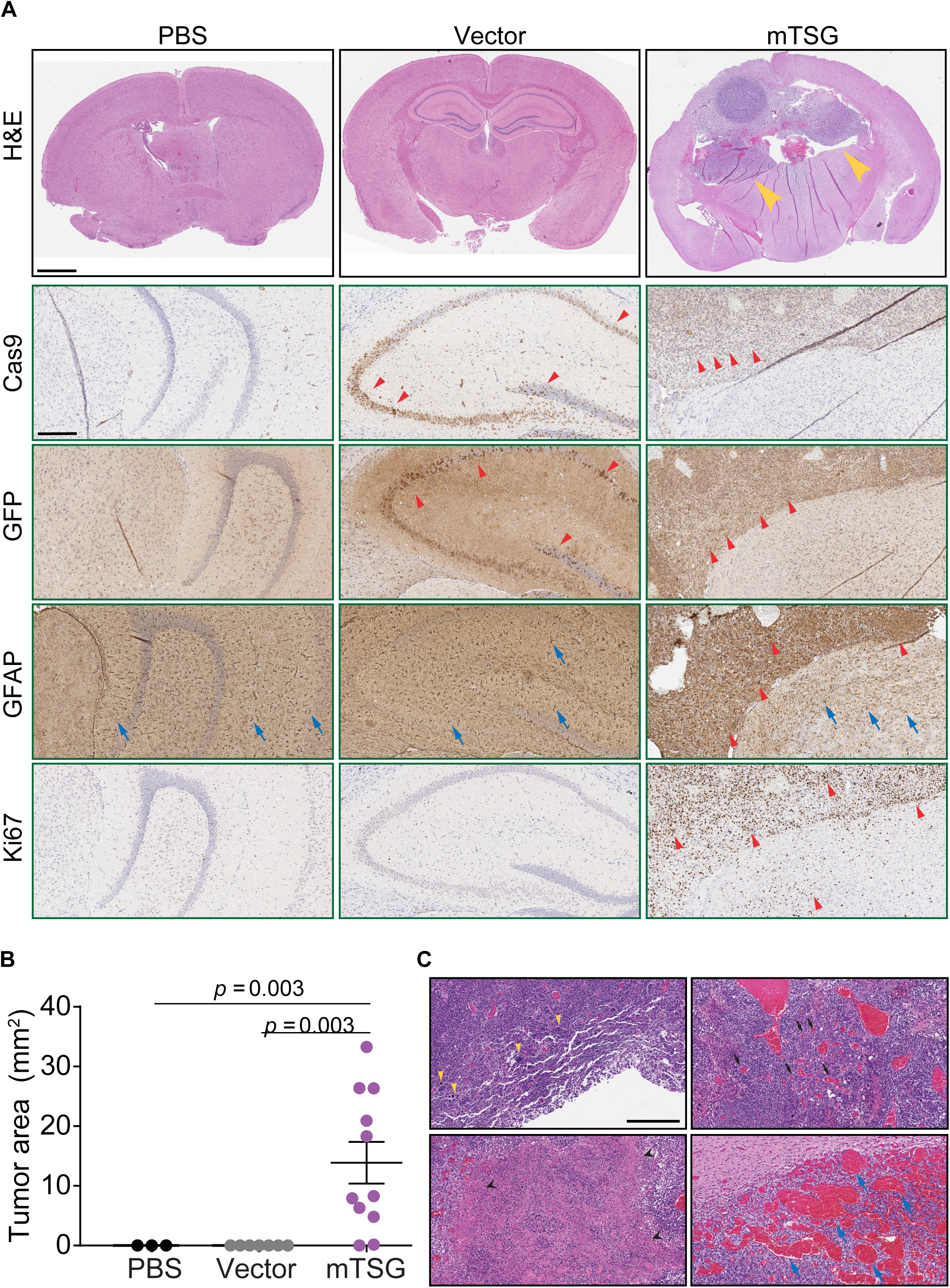
Histopathological analysis of AAV-mTSG induced brain tumors revealed pathological features of GBM. **A.** (Top) Representative images of endpoint histology (H&E) brain sections from PBS, AAV-vector and AAV-mTSG injected mice. Arrowhead indicates brain tumor. Scale bar = 1 mm. (Lower) Representative images of immunostained brain sections from PBS, AAV-vector and AAV-mTSG injected mice. Cas9 IHC, arrowheads indicate Cas9-positive cells in the injected brain regions (AAV-vector) and tumors (AAV-mTSG) but not non-tumor regions; GFP IHC, arrowheads indicate GFP-positive cells in the injected brain regions (AAV-vector) and tumors (AAV-mTSG) but not non-tumor regions; GFAP IHC, representative GFAP-positive astrocytes in PBS, AAV-vector and AAV-mTSG injected mice (blue arrows), as well as representative cancer cells in AAV-mTSG injected mice (read arrowheads); Ki67 IHC, arrowheads indicate representative proliferative cells, which are mostly in tumors (AAV-mTSG) or scattered in tumor-adjacent brain regions (AAV-mTSG). Scale bar = 0.25 mm. **B.** Quantification of tumor sizes found in H&E brain sections from PBS, AAV-vector and AAV-mTSG injected mice. Two-tailed t-test, *p* = 0.003, mTSG vs. vector or PBS (PBS, n = 3; Vector, n = 7; mTSG, n = 11). **C.** Representative higher magnification H&E images showing pathological features of AAV-mTSG induced mouse GBM. Left to right, yellow arrowheads in the first panel indicate representative giant aneuploid cells with pleomorphic nuclei; Black arrows in the second panel indicate representative endothelial cells and angiogenesis; Black arrowheads in the third panel indicate representative necrotic regions; Blue arrows in the fourth panel indicate representative hemorrhage regions. Similar features were observed in histology analysis of human GBM patient sections from Yale Glioma tissue bank. Scale bar = 0.5 mm.

Because AAVs usually do not integrate into the genome, direct sequencing of the targeted regions was needed to determine which mutations (and genes associated with them) were in each tumor. To globally map the molecular landscape of these brain tumors, we designed a customized probe set (mTSG-Amplicon probes) covering the target regions of all sgRNAs in the mouse genome (Methods) (Table S5). We used these probes to perform targeted captured sequencing for whole brain samples and liver samples (as a control organ not being directly transduced) in a cohort of AAV-mTSG, AAV-vector and PBS injected mice (n = 25, 3, and 4, respectively) (Table S6). As a result, 277/278 (99.6%) of unique sgRNA target regions were captured for all samples from this experiment, with the missing one being Arid1a-sg5 (due to unavailability of qualified regions in capture probe design). Across all 40 samples, the average of mean coverage across all regions flanking sgRNA cutting sites was 19,405 ± 180 (Table S7). We analyzed the mutant variants at the predicted cutting sites of the 277 successfully captured unique sgRNAs across all samples (Methods) (Table S8). At the single sgRNA level, for example, at the predicted cutting site of sgRNA-4 in the *Mll2* locus (also known as *Kmt2d*), various insertions and deletions (indels) were detected in AAV-mTSG but not AAV-vector or PBS mice (Figure 3A). As gliomagenesis takes multiple months in mice (Figure 1D), we surveyed 3 mice at 3.5 weeks post-injection and performed capture sequencing to revealed early mutation profiles, as an approximation for *in vivo* sgRNA cutting efficiency (Figure S4A-B, Table S9). We found that even low-efficiency sgRNAs can end up being highly enriched in the process of tumorigenesis if the mutations they generated are strongly oncogenic (Figure S4C, Figure 3B-C). After removing represent germline variants, we determined whether the regions flanking each sgRNA target site would be classified as significantly mutated sgRNA sites (SMSs) (Methods) (Table S10). Specifically, we implemented a false-discovery-rate (FDR) approach with (FDR < 1/12, or 8%) as well as a flat 5% variant frequency cutoff, and confirmed that the choice of alternative cutoffs did not alter the final SMS calls (Figure S5B). With these criteria, we observed a diverse mutational landscape across most mice that were capture-sequenced (Figure 3B-C, Figure S5A, Table S11). As an example, one AAV-mTSG mouse (mTSG brain 25) had significant mutations at the predicted cutting sites of 16 out of 277 captured gene-targeting sgRNAs in the mTSG library, covering 12 significantly mutated genes (mSMGs) (Figure 3B). A second example (mTSG brain 39) showed a more diverse mutational profile (34 SMSs for 26 mSMGs) (Figure 3B). The raw indel frequencies were also summed across all detected variants for each sgRNA target site in each sample, revealing a highly diverse pattern of variant frequencies generated by this sgRNA pool (Figure 3C) (Table S10). Comparing brain samples between treatment groups, AAV-mTSG injected brains had significantly higher mean variant frequencies (2.087 ± 0.429, n = 25) compared to vector (0.005 ± 0.001, n = 3) or PBS (0.003 ± 0.001, n = 4) injected brains (two-tailed Welch’s t-test, *p* < 0.0001 for both mTSG vs. vector and mTSG vs. PBS) (Figure 3C-D). Comparing targeted vs. non-targeted organs in AAV-mTSG injected mice, the mean variant frequencies of brains (2.087 ± 0.429, n = 25) were significantly higher than livers (0.309 ± 0.261, n = 4) (two-tailed Welch’s t-test, *p* = 0.002) (Figure 3C-D). As shown in a meta-analysis of all variants of all sgRNAs for each sample, the predominant indels were deletions for virtually all samples, and most insertions at SMS sites were 1 bp in size (Figure 3E) (Table S8). We identified distinct variant frequency clusters of sgRNA-induced indels that may serve as an approximation to the clonality of these tumors (Methods). From this analysis, we found that only 2/22 of the brains had monoclonal tumors, with the majority (20/22) being comprised of multiple clusters (Figure S5C). These data demonstrate on-target, pooled genome editing in the brain at a library scale, stochastically generating loss-of-function mutations in native glial cells and priming driving them for subsequent selection during gliomagenesis.

**Figure 3.**
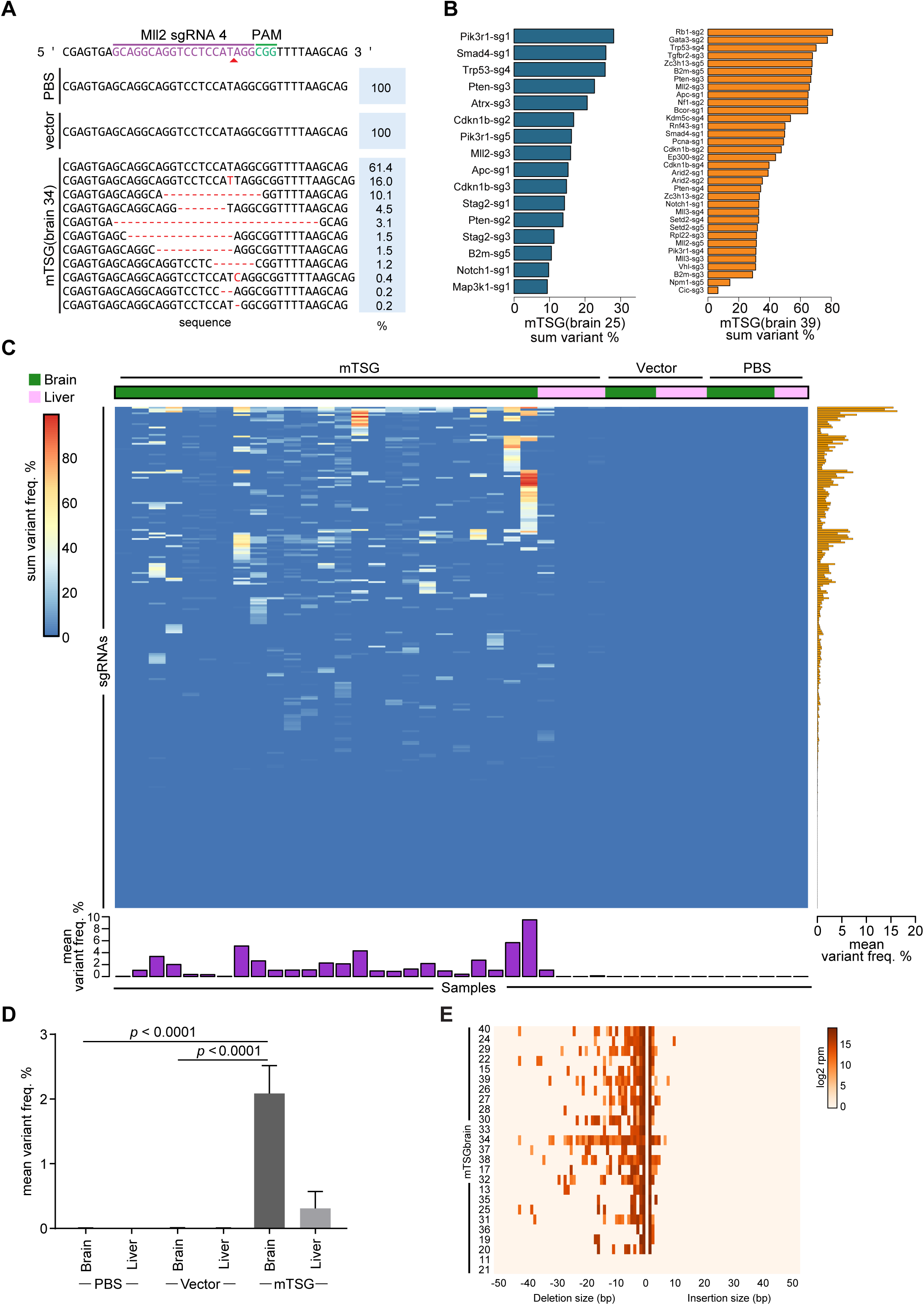
Targeted captured sequencing of sgRNA sites showed a global variant map in AAVmTSG induced mouse GBM. **A.** Representative indel variants generated by individual sgRNA target regions. Alleles observed at the genomic region targeted by *Mll2* sgRNA 4 in representative PBS, AAV-vector, and AAV-mTSG injected mouse brain samples. The percentage of total reads that correspond to each allele is indicated on the right (blue box). **B.** Representative mutation profiles of all sgRNA target regions in individual samples. Bar plots showing two representative AAV-mTSG injected mouse brain samples with variant frequency in significantly mutated sgRNA target regions. Sum variant frequency is the cumulative frequency of all detected variants for a particular sgRNA. **C.** A global heatmap of sum indel frequency across all targeted capture samples. Each row represents the sum indel frequencies of one sgRNA across samples. Each column is a brain sample from mice stereotaxically injected with PBS, AAV-vector, or AAV-mTSG. Liver was a non-targeting organ and thus was used as a background control. Most sgRNAs in most samples have low mean variant frequency as shown in dark blue color. SgRNAs with higher mean variant frequencies were evident in a subset of mTSG samples but not vector or PBS samples. One exception is *Rps19* sgRNA 5, which had a moderate variant frequency in all samples, representing germline variants in LSL-Cas9 mice, and was thus excluded. Bar plots of the mean average variant frequencies for each sgRNA (right panel, orange bars) and each sample (bottom panel, purple bars) are also shown. **D.** Barplot of mean variant frequency, grouped by treatment condition and tissue type. AAV-mTSG injected brains had significantly higher mean variant frequencies (2.087 ± 0.429, n = 25) compared to vector (0.005 ± 0.001, n = 3) or PBS (0.003 ± 0.001, n = 4) injected brains (two-tailed t-test, *p* < 0.0001 for both mTSG vs. vector and mTSG vs. PBS). Comparing targeted vs. non-targeted organs in AAV-mTSG injected mice, mean variant frequencies of brains (2.087 ± 0.429, n = 25) were significantly higher than livers (0.309 ± 0.261, n = 4) (two-tailed t-test, p = 0.002) **E.** Metaplot heatmap of indel size distribution for all filtered variants in each of the mTSG brain samples (rows, n = 25). A negative number of basepairs indicates deletions; positive numbers of basepairs indicate insertions; color as in key (left subpanel) indicate the relative abundance of indels of a particular size in a particular mouse.

We next summarized the mutational data from the SMS level to the mSMG level (Methods) (Table S12) and created an oncomap of all mTSG brain samples (Figure 4A). Across all AAV-mTSG samples, the number of SMSs ranged from 3 to 46, with mSMGs ranging from 3 to 33 (Table S11). Across all mice with SMGs (23/25), the detected variants were predominantly frameshift indels (frameshift reads / total variant fraction >60% in 22/23 mice) compared to non-frameshift indels, splicing indels and intronic indels (Figure 4A, bottom panel). Surprisingly, all 56 genes have at least one associated SMSs, with eight of them (*Pten, Map2k4, B2m, Pcna, Cic, Setd2, Gata3* and *Apc*) having 5/5 SMSs (Figure 4A left and middle panel). The mSMGs encompass functionally diverse categories of proteins, including cell death or cell cycle regulators, immunological regulator, DNA repair and replication regulators, transcriptional repressors, epigenetic regulators, transcription factors, cadherin type proteins and ubiquitin ligases (Figure 4A). Many of the mSMGs are significantly mutated in 20% to 50% of mice, with most of the epigenetic regulators in this range, such as *Arid1b, Mll3, Setd2, Mll2, Kdm5c, Kdm6a, Arid2* and *Ctcf* (Figure 4A), highlighting the central importance of epigenetic regulators in tumorigenesis in the brain. Interestingly, *B2m*, a core component of major histocompatibility complex (MHC) class I that is essential for antigen presentation, appeared as the second most frequently mutated gene (19/25 = 76% of mice) (Figure 4A, right panel). This pooled direct mutagenesis analysis revealed, in a quantitative manner, the relative phenotypic strength of loss-of-function mutations in the most significantly mutated genes, for driving gliomagenesis *in vivo*.

**Figure 4.**
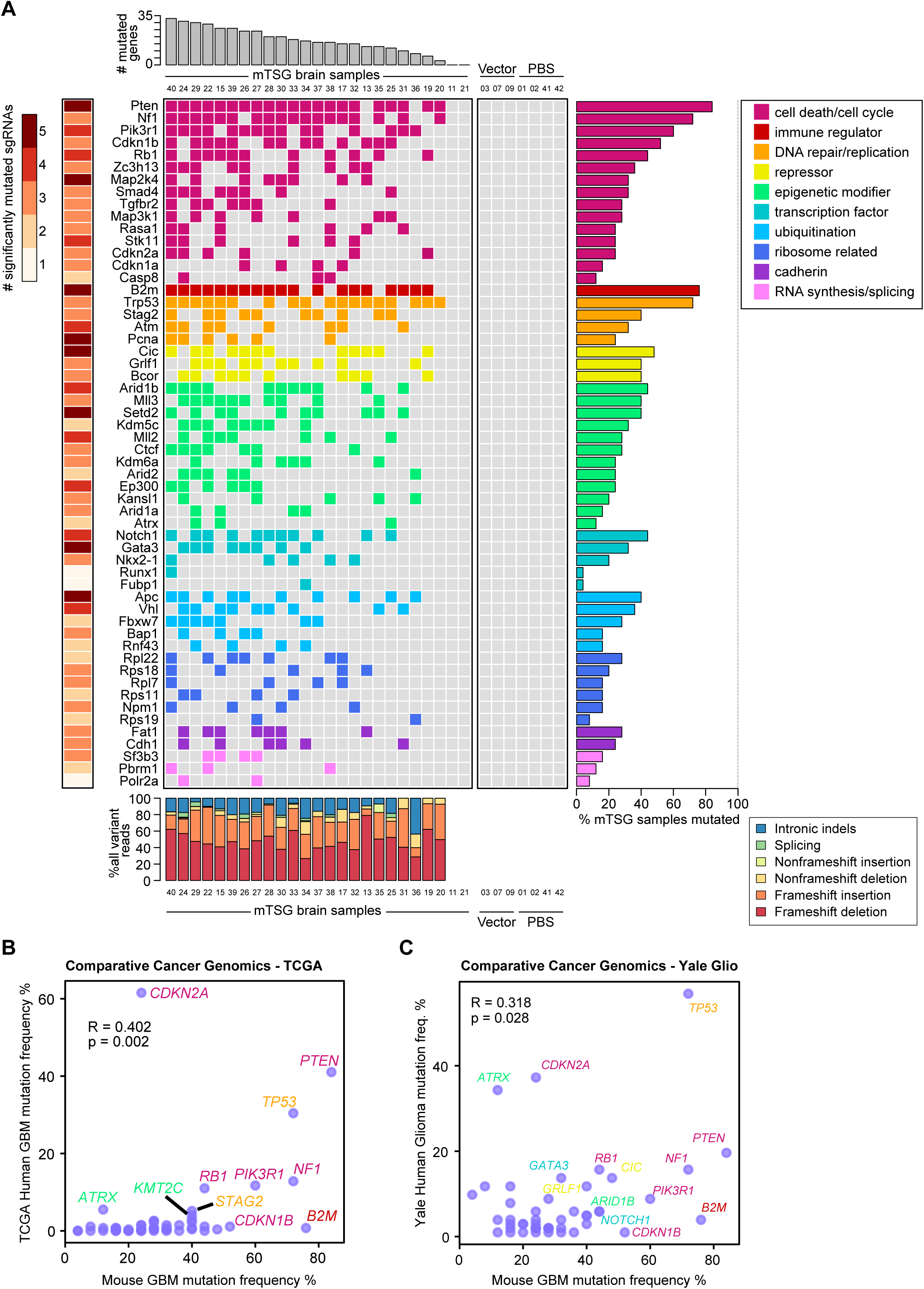
Integrative and comparative cancer genomic analysis of functional SMGs in gliomagenesis. **A. Gene-level mutational landscape of AAV-mTSG induced primary mouse GBM** **Center:** Tile chart depicting the mutational landscape of primary brain samples from LSL-Cas9 mice infected with the AAV-mTSG library (n = 25), AAV-vector (n = 3) or PBS (n = 4). Genes are grouped and colored according to their functional classifications (cell death/cycle, immune regulator, DNA repair/replication, repressor, epigenetic modifier, transcription factor, ubiquitination, ribosome related, cadherin, and RNA synthesis/splicing), as noted in the legend in the top-right corner. Colored boxes indicate that the gene was significantly mutated in a given sample, while a gray box indicates no significant mutation. **Top:** Bar plots of the total number of significantly mutated genes identified in each AAV-mTSG sample, labeled with sample number in the bottom (n = 25). **Right:** Bar plots of the percentage of GBM samples that were called as significantly mutated for each gene. *Pten, B2m, Trp53, Cic, Arid1b, Notch1, Apc, Rpl22, Fat1* and *Sf3b3* were the top mutated genes in each of the 10 functional classifications, respectively. **Left:** Heatmap of the numbers of unique significantly mutated sgRNAs (SMSs) for each gene. **Bottom:** Stacked bar plots describing the type of indels observed in each sample, including intronic indel, splice-site indel, non-frameshift insertion, non-frameshift deletion, frameshift insertion and frameshift deletion, color-coded according to the legend in the bottom-right corner. **B.** Comparative cancer genomics in GBM using TCGA dataset. Scatterplot of gene population-wide mutant frequencies for the genes represented in the mTSG library, comparing AAV-mTSG treated mouse brain samples to human samples. Pearson correlation coefficient is shown on the plot, revealing that the mutation frequencies in this multiplexed mutagenesis mouse model were significantly correlated with those in human patients (R = 0.402, *p* = 0.002). Representative strong drivers in both species were shown, with gene names color-coded based on their functional classification (as in Figure 4), showing predominant drivers in the categories of cell death / cell cycle (*PTEN, NF1, CDKN2A, PIK3R1, RB1, CDKN1B*), DNA repair / replication (*TP53, STAG2*), immune regulator (*B2M*) and epigenetic modifier (*ATRX, KMT2C/MLL3*). **C.** Comparative cancer genomics in GBM using Yale Glioma dataset (Yale Glio). Scatterplot of gene population-wide mutant frequencies for the genes represented in the mTSG library, comparing AAV-mTSG treated mouse brain samples to human samples. Pearson correlation coefficient is shown on the plot, revealing that the mutation frequencies in this multiplexed mutagenesis mouse model were significantly correlated with those in human patients (R = 0.318, *p* = 0.028). Representative strong drivers in both species were shown, with gene names color-coded based on their functional classification (as in Figure 4), showing predominant drivers in the categories of cell death / cell cycle (*PTEN, NF1, CDKN2A, PIK3R1, RB1, CDKN1B*), DNA repair / replication (*TP53*), immune regulator (*B2M*), epigenetic modifier (*ATRX, ARID1B*), repressor (*CIC, GRLF1*) and transcription factor (*NOTCH1, GATA3*).

We compared the mutational frequencies in mice to the variant frequencies of their homologous genes in human GBM with their frequencies of non-silent mutation and deletion (Methods). For these 56 genes, the mutation frequencies in mouse GBMs (an end-product of pooled mutagenesis and *in vivo* gliomagenesis) significantly correlated with the mutation frequencies in TCGA GBM patients (Pearson correlation coefficient R = 0.402, *p* = 2.1 * 10^−3^) (Figure 4B) (Table S13). An outlier was *Cdkn2a*, which was mutated at higher frequencies in human compared to those in mice (outlier test, p < 0.05) (Figure 4B). To further investigate this correlation, we utilized the clinical cancer genomics data from the Yale Glioma tissue bank, a source independent from TCGA (Tables S14, S15). Collectively, the mouse mutation frequencies again significantly correlated with those in human patients (R = 0.318, *p* = 0.0277) (Figure 4C) (Table S16). Summarizing from the mutation frequencies in mouse and both human GBM patient cohorts, the strongest tumor suppressors in both species belong to the functional categories of cell death / cell cycle regulator (*PTEN, NF1, CDKN2A, PIK3R1, RB1, CDKN1B*), DNA repair / replication regulator (*TP53, STAG2*), immune regulator (*B2M*), repressor (*CIC, GRLF1*), transcription factor (*NOTCH1, GATA3*) and epigenetic regulator (*ATRX, KMT2C/MLL3, ARID1B*) (Figure 4B-C). These data suggest that the AAV-CRISPR library-based autochthonous GBM mouse model revealed a quantitative phenotypic profile of tumor suppressors reflecting the genomic landscape of human GBM patients.

To generate an unbiased map of co-drivers, we calculated the co-occurrence rate and the statistical significance of double-mutations for each pair of genes (Methods) (Figure 5A-B, Table S17). This analysis showed that 76 gene pairs out of a total of 1540 all possible pairs were statistically significant in terms of co-occurrence (hypergeometric test *p* < 0.0025, FDR adjusted *q* < 0.05). The *Nf1*/*Pten* pair emerged as the top pair by the strength of co-occurrence (co-occurrence rate = 18/21 = 85.7%, hypergeometric test, *p* = 7.53 * 10^−8^) (Figure 5A-C), consistent with this gene pair functioning as strong tumor suppressors of GBM in humans. Interestingly, several previously undocumented combinations emerged with high rates of co-occurrence and statistical significance, such as such as *Kdm5c*/*Gata3* (co-occurrence rate = 77.8%, hypergeometric test, *p* = 6.04 * 10^−6^), and *B2m/Pik3r1* (70.0%, *p* = 2.28 * 10^−5^) (Figure 5A-C). In addition, we performed correlation analysis of summed mutant frequencies for each pair of genes across all individual mice (Methods) (Figure 5D-E, Table S18). 22.9% (352/1540) of the gene pairs were significantly positively correlated (Spearman correlation > 0, *p* < 0.0116, FDR adjusted *q* < 0.05) (Figure 5D-E). The most significantly correlated gene pair was again *Nf1/Pten* (Spearman correlation = 0.861, *p* = 3.34 * 10^−8^) (Figure 5D-F, Table S18), along with other representative pairs such as *Cdkn2a/Ctcf* (correlation = 0.792, *p* = 2.41 * 10^−6^) (Figure 5G), *B2m/Notch1* (correlation = 0.789, *p* = 2.82 * 10^−6^), and *Apc/Pik3r1* (correlation = 0.774, *p* = 5.77 * 10^−6^) (Figure 5D-E). Exclusion of the master sgRNA against *Trp53* revealed largely identical results for the remaining genes (Figure S6A-B). Of note, a subset of the significantly co-occurring pairs were also found to be co-mutated in human GBM, including the pairs of *RB1*+*TP53*, *PTEN*+*RB1*, *RASA1*+*STK11*, *B2M*+*MAP2K4*, *PTEN*+*STAG2*, *CDKN1B*+*TP53* and *CDKN1B*+*NF1* (Figure S6C-D). These data revealed systematic co-occurrence and correlation relationships of specific mutations during glioblastoma progression *in vivo*.

**Figure 5.**
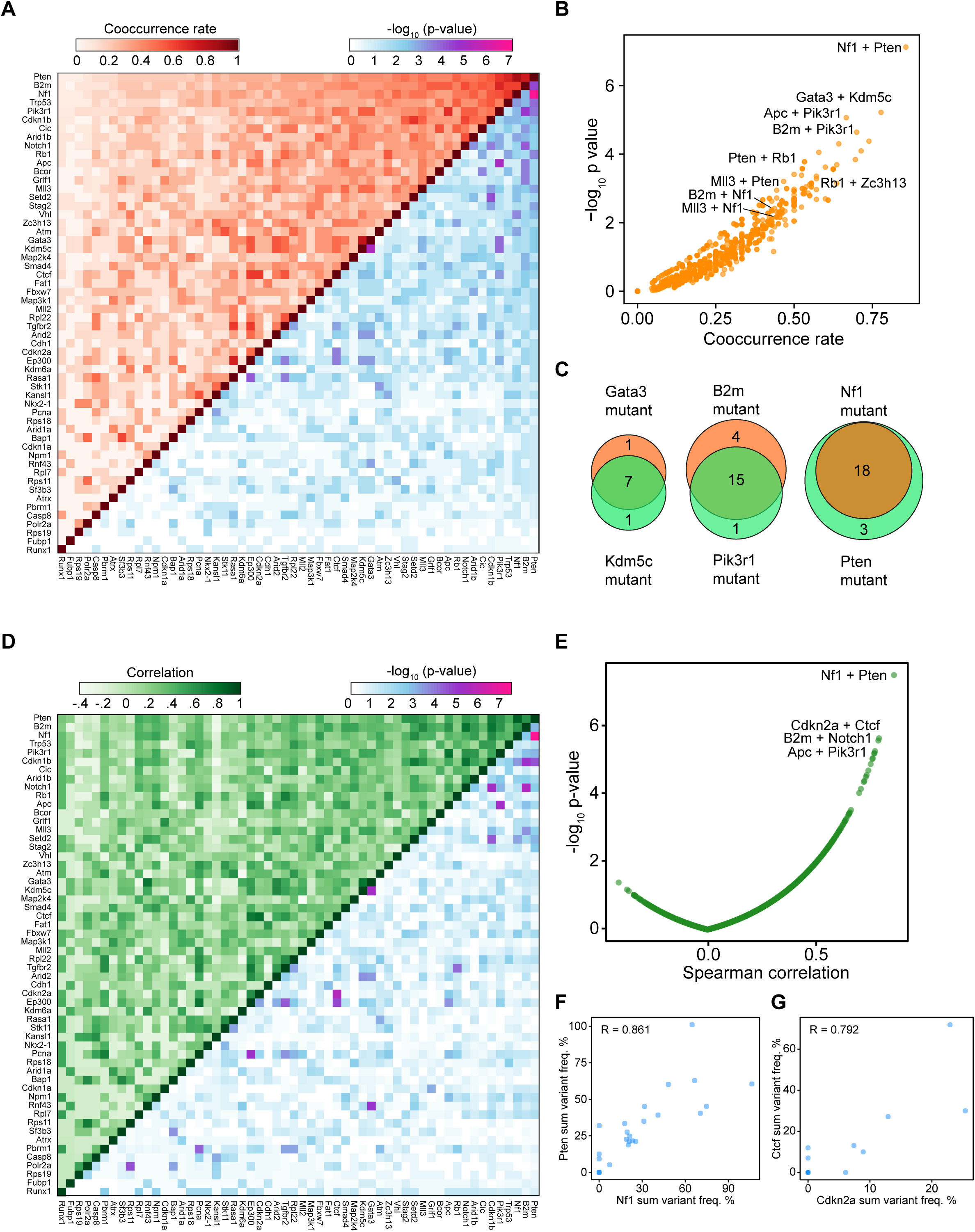
Co-occurrence and correlation analyses of GBM mSMGs uncovered synergistic gene pairs. **A.** Upper-left half: heatmap of the number of samples with a mutation in both specified genes. Lower-right half: heatmap of −log_10_ p-values by hypergeometric test to evaluate whether specific pairs of genes are statistically significantly co-mutated. Co-occurrence rate of each pair is defined by the intersection over union, where the intersection is defined as the number of double-mutant samples, and the union as the number of samples with a mutation in either of the two genes. **B.** Scatterplot of the cooccurrence rate of a given gene pair, plotted against −log_10_ p-values. Representative co-occurring pairs are indicated. **C.** Venn diagrams showing representative strongly co-occurring mutated gene pairs such as *Kdm5c* and *Gata3* (co-occurrence rate = 77.8%, hypergeometric test, *p* = 6.04 * 10^−6^), *B2m* and *Pik3r1* (70.0%, *p* = 2.28 * 10^−5^), as well as *Nf1* and *Pten* (85.7%, *p* = 7.53 * 10^−8^). **D.** Upper-left half: heatmap of the pairwise Spearman correlation of sum variant frequency for each gene, averaged across sgRNAs. Lower-right half: heatmap of −log_10_ p-values to evaluate the statistical significance of the pairwise correlations. **E.** Scatterplot of pairwise Spearman correlations plotted against −log_10_ p-values. Representative top pairs are indicated. **F-G.** Scatterplots showing representative strongly correlated gene pairs when comparing sum % variant frequencies averaged across sgRNAs, such as *Nf1*+*Pten* (**F)** and *Cdkn2a*+*Ctcf* (**G)**. The Spearman correlation coefficients are noted on the plot.

We went on to test several of the highly represented individual drivers or combinations using an sgRNA minipool validation approach (Methods) (Figure 6A). All of the uninjected (n = 2), EYFP (n = 4) and empty vector (n = 3) injected mice survival and maintain good body condition for the whole duration of the study, and were devoid of any observable tumors by histology analysis with mice sacrificed from 4 to 11 months after treatment (Figure 6B, Figure S7, Table S19), again indicating that without mutagenesis, or with *Trp53* disruption alone, LSL-Cas9 mice did not develop brain tumors. In contrast, within 11 months post injection, 50% (4/8) of mice receiving AAVs containing *Nf1* sgRNA minipool developed macrocephaly, poor body condition score and large tumors (compared to all 9 control mice, two-tailed Fisher’s exact test, *p* = 0.029). All mice receiving *Nf1*;*Pten* (9/9, 100%, two tailed Fisher’s exact test, *p* = 4.11 *10^−5^) and *Nf1*;*B2m* minipools (4/4, 100%, *p* = 0.0014) developed macrocephaly, poor body condition score and large tumors (Figure 6C, Table S19). All mice receiving *Rb1, Rb1;Pten,* or *Rb1;Zc3h13* minipools (3/3, 100%, *p* = 0.0045 for all three groups) developed macrocephaly, poor body condition score and large tumors (Figure 6D, Table S19). For the same duration of study (maximum 11 months), smaller fractions of mice receiving the AAV sgRNA minipools targeting *Arid1b;Nf1* (4/9), *Mll3;Nf1* (2/5), *Mll2* (2/10), *Cic* (1/5), *Cic;Pten* (1/4), *Setd2* (1/5), and *Gata3;Mll3* (1/5) (Table S19). Collectively, half (40/80, or 50%) of the mice receiving AAV sgRNA minipools targeting any of the single gene or gene pairs developed brain tumors within 11 months (Collective validation vs. all controls, two-tailed Fisher’s exact test, *p* = 0.004). These data indicated that mutating these individual genes or combinations in combination with *Trp53* causes GBM in fully immunocompetent animals.

**Figure 6.**
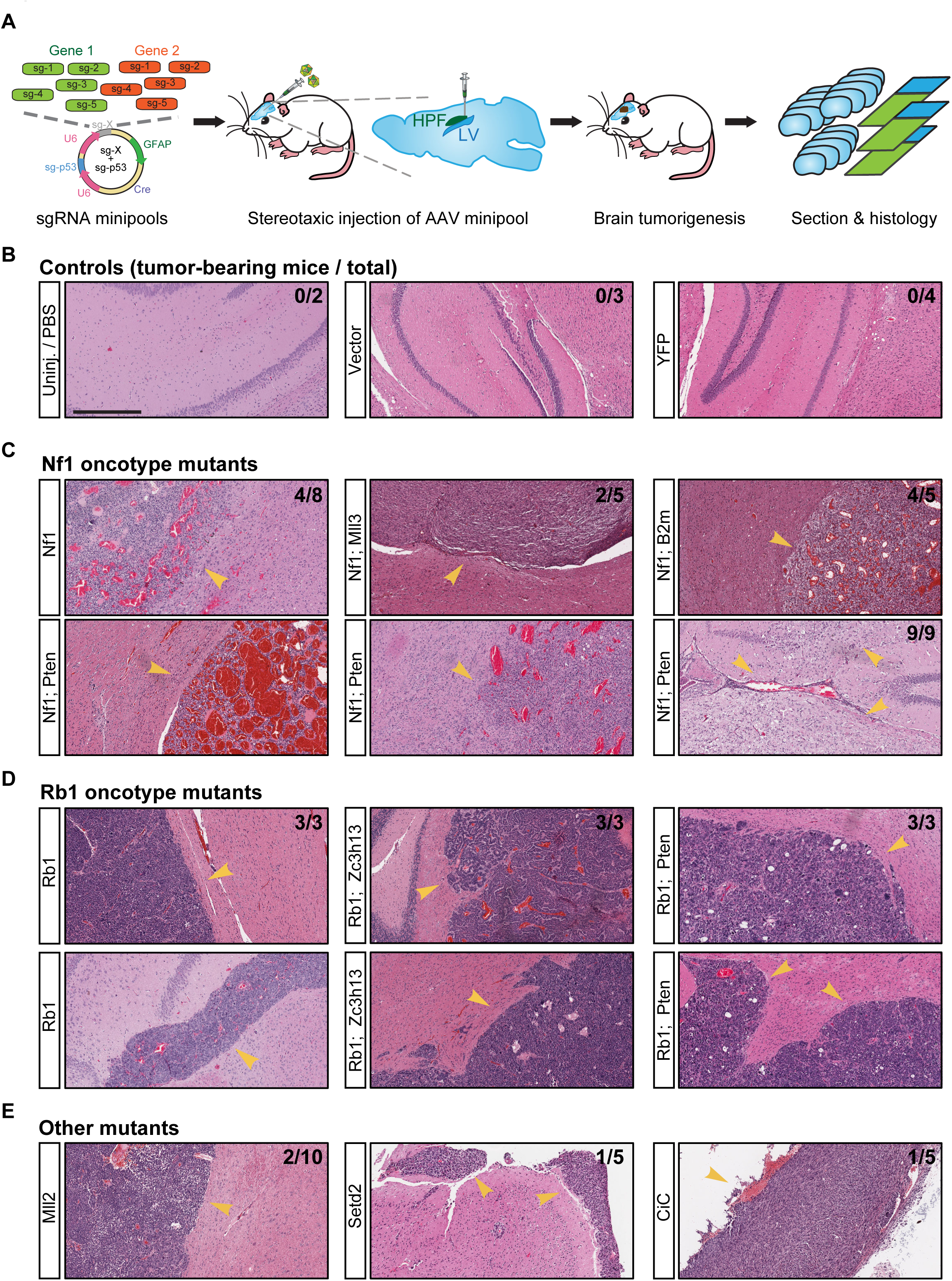
Validation of driver combinations. **A.** Schematic representation of experiment design. Mixtures of five sgRNAs targeting each gene were cloned as sgRNA minipool into the same astrocyte-specific AAV-CRISPR vector. After packaging, AAV minipools were stereotaxically injecting into the lateral ventricle of LSL-Cas9 mice. Mice were euthanized when either macrocephaly or poor body condition score was observed, and brains were isolated to perform sectioning and end-point histology analysis. **B-E.** End-point histology (H&E) of representative brain sections from mice treated with AAV sgRNA minipools or relevant controls. In this end-point analysis, mice were euthanized when either macrocephaly or poor body condition score (< 2) was observed, with survival time ranging from 3 to 11 months. Treatments are indicated in the leftmost boxes. Arrowheads indicate the presence of brain tumors in mice receiving the sgRNA AAV minipools. The proportion of tumor-bearing to total mice is indicated in the top right corner of the images. Scale bar = 0.5 mm. **B.** Representative histology of brain sections from control mice. No tumors were observed in mice from the vector (0/3), EYFP (0/4) or uninjected (0/2) groups, all of which maintained good body condition without macrocephaly for the entire duration of the study. Control mice were euthanized for brain sectioning and histology analysis at 4 to 11 months post injection. **C.** Representative histology of brain sections from mice treated with various *Nf1* minipools, such as *Nf1* alone (4/8 mice developed tumors within 11 months), *Nf1;Pten* (9/9 mice developed tumors within 6 months), *Nf1;Mll3* (2/5 mice developed tumors within 6 months), and *Nf1;B2m* (4/5 mice developed tumors within 11 months). **D.** Representative histology of brain sections from mice treated with various *Rb1* minipools, such as *Rb1* alone (3/3 mice developed tumors within 6 months), *Rb1;Zc3h13* (3/3 mice developed tumors within 6 months), and *Rb1;Pten* (3/3 mice developed tumors within 6 months). **E.** Representative histology of brain sections from mice treated with other minipools, such as *Mll2* alone (2/10 mice developed tumors within 11 months), *Setd2* (1/5 mice developed tumors within 6 months), and *Cic* (1/5 mice developed tumors within 6 months).

Interestingly, brain tumors with *Nf1* mutations displayed highly polymorphic pathological features, with diverse, fibroblastic cell morphologies, frequently with regions of necrosis, and large-area hemorrhage (Figure 6C), and are almost always GFAP-positive (Figure S7), whereas in sharp contrast, tumors with *Rb1* mutations composed of round cells with dense nuclei, frequently with proliferative spindles and giant cells with massive nuclear aneuploidy and pleiomorphism, but rarely with regions of necrosis or large-area hemorrhage (Figure 6D), and often contain mixtures of GFAP-positive and GFAP-negative cells (Figure S7). Intrigued by the morphologically distinct pathology of *Nf1* and *Rb1* mutant tumors, as well as the newly validated, but previously undocumented driver combinations, we went on to investigate the molecular regulation of gliomagenesis driven by different combinations of drivers (Methods) (Figure 7A, Figure S8A). To gain a global quantitative picture of the molecular changes underlying these different mutations, we performed mRNA-Seq to profile the transcriptome of these mutant glioma cells (*Nf1, Nf1;Mll3, Rb1,* and *Rb1;Zc3h13,* n = 3 biological replicates each) (Tables S20, S21). Comparing *Nf1* mutant and *Rb1* mutant cells, we found that 616 genes were more highly expressed in *Rb1* cells (Benjamini-Hochberg adjusted *p* < 0.05 and log fold change ≥ 1), while 982 genes were more highly expressed in *Nf1* cells (adjusted *p* < 0.05 and log fold change ≤ −1) (Figure 7B, Table S22). Gene ontology analysis of the genes associated with higher expression in *Nf1* mutant cells revealed multiple enriched categories (Benjamini-Hochberg adjusted *p* < 0.05), including extracellular region part, biological adhesion, cell adhesion, neuron differentiation, extracellular matrix, hormone metabolic process, cell motion, and cell-cell signaling (Figure 7C). Gene ontology analysis of the genes associated with higher expression in *Rb1* mutant cells revealed a distinct set of enriched categories (adjusted *p* < 0.05), which surprisingly included regionalization, anterior/posterior pattern formation, transcription factor activity, embryonic morphogenesis, cell adhesion, extracellular matrix, neuron differentiation, and GTPase regulator activity (Figure 7D). Strikingly, a total of 13 *Homeobox* genes (*Hoxb9, Hoxb8, Hoxa11, Hoxc10, Hoxb3, Hoxa9, Hoxb7, Hoxa7, Hoxb2, Hoxc11, Hoxa10, Hoxb6* and *Hoxb5*) were among the top-40 upregulated genes in *Rb1* mutants. The enrichment of multiple development-related gene ontologies in *Rb1* mutant cells vs. *Nf1* mutant cells thus reflects a molecular alteration of differentiation status in *Rb1* tumors. Consistent with this notion, we observed that *Rb1* mutant tumors exhibit histological characteristics that are suggestive of a lineage transformation event away from astrocytes: as previously noted, *Rb1* tumors were composed of round cells with large dense nuclei that comprised the vast majority of the total cell size (Figure 6D), and often with mixtures of GFAP-positive and GFAP-negative cells (Figure S7).

**Figure 7.**
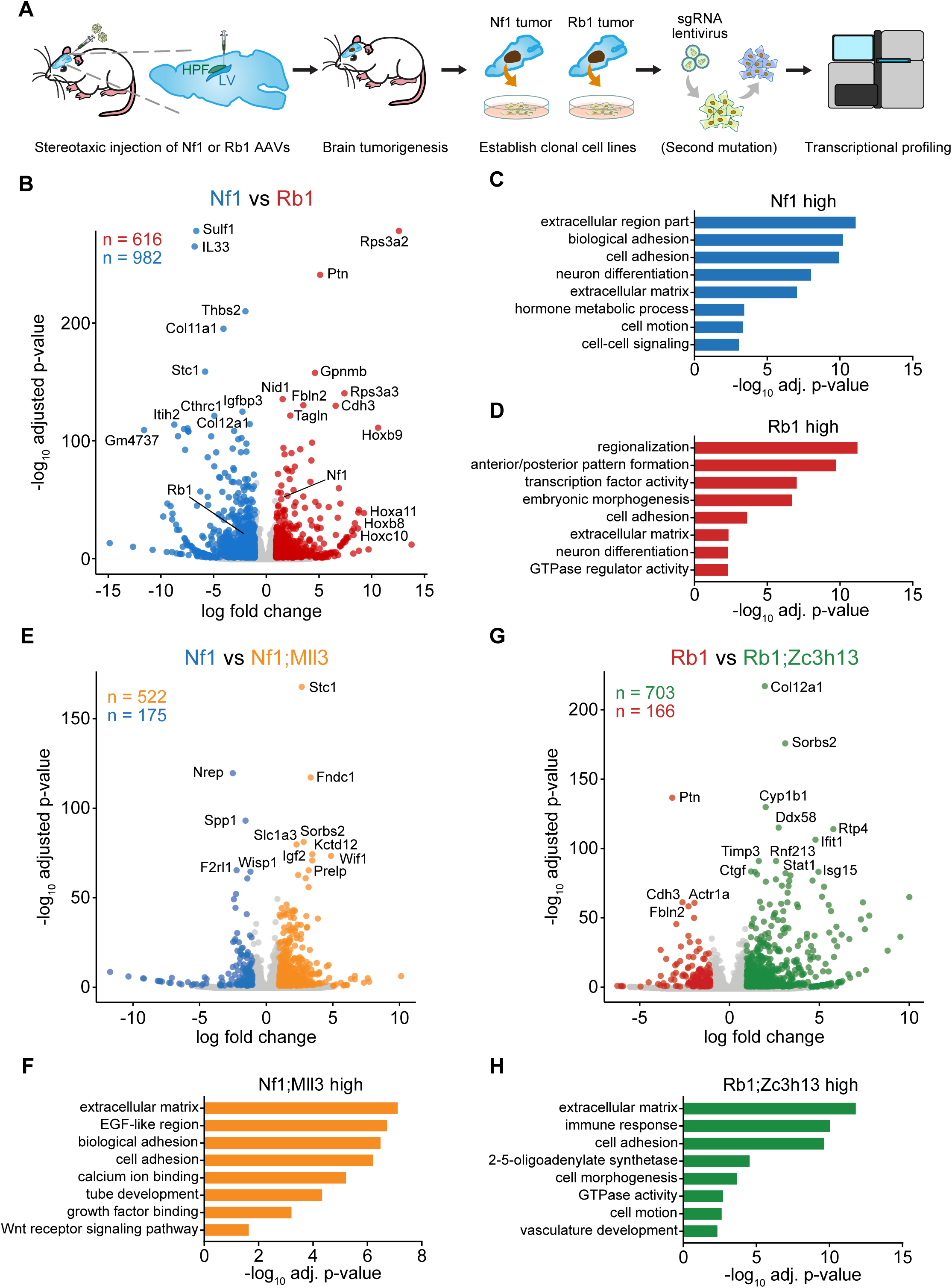
Transcriptome profiling of mouse GBM driver combinations. **A.** Schematic of mouse GBM mutant RNAseq experimental design. *Rb1* or *Nf1* AAV minipools were stereotaxically injected into the lateral ventricle of LSL-Cas9 mice. Cell lines were derived from mouse GBMs by single-cell isolation, plating and culture in DMEM media. Additional driver mutations were introduced by lentiCRISPR where applicable. Cells were harvested for transcriptome profiling by RNA-Seq. **B.** Volcano plot comparing *Rb1* mutant (red) to *Nf1* mutant (blue) GBM cells. 616 genes were significantly higher in *Rb1* cells (Benjamini-Hochberg adjusted *p* < 0.05 and log fold change ≥ 1), and 982 genes were significantly higher in *Nf1* cells (adjusted *p* < 0.05 and log fold change ≤ −1). Examples of highly differentially expressed genes were labeled. Noted as an on-target benchmark, *Nf1* was expressed at a significantly lower level in *Nf1* mutant cells, and *Rb1* was expressed at a significantly lower level in *Rb1* mutant cells. **C.** Enriched gene ontology categories among *Nf1*-high genes. Top categories included extracellular region part, biological adhesion, cell adhesion, neuron differentiation, extracellular matrix, hormone metabolic process, cell motion, and cell-cell signaling. Negative log_10_ Benjamini-Hochberg adjusted p-values are shown. **D.** Enriched gene ontology categories among *Rb1*-high genes. Top categories included regionalization, anterior/posterior pattern formation, transcription factor activity, embryonic morphogenesis, cell adhesion, extracellular matrix, neuron differentiation, and GTPase regulator activity. Negative log_10_ Benjamini-Hochberg adjusted p-values are shown. **E.** Volcano plot comparing *Nf1;Mll3* mutant (orange) to *Nf1* mutant (blue) GBM cells. 522 genes were significantly higher in *Nf1;Mll3* cells (Benjamini-Hochberg adjusted *p* < 0.05 and log fold change ≥ 1), and 175 genes were significantly higher in *Nf1* cells (adjusted *p* < 0.05 and log fold change ≤ −1). Examples of highly differentially expressed genes were labeled. **F.** Enriched gene ontology categories among *Nf1;Mll3*-high genes. Top categories included extracellular matrix, EGF-like region, biological adhesion, cell adhesion, calcium ion binding, tube development, growth factor binding, and Wnt receptor signaling pathway. Negative log_10_ Benjamini-Hochberg adjusted p-values are shown. **G.** Volcano plot comparing *Rb1;Zc3h13* mutant (green) to *Rb1* mutant (red) GBM cells. 703 genes were significantly higher in *Rb1;Zc3h13* cells (Benjamini-Hochberg adjusted *p* < 0.05 and log fold change ≥ 1), and 166 genes were significantly higher in *Rb1* cells (adjusted *p* < 0.05 and log fold change ≤ −1). Examples of highly differentially expressed genes were labeled. **H.** Enriched gene ontology categories among *Rb1;Zc3h13*-high genes. Top categories included extracellular matrix, immune response, cell adhesion, 2-5-oligoadenylate synthetase, cell morphogenesis, GTPase activity, cell motion, and vasculature development. Negative log_10_ Benjamini-Hochberg adjusted p-values are shown.

*Mll3* encodes a lysine N-methyltransferase ^32^. *Zc3h13* is a largely uncharacterized zinc-finger protein that has been shown to interact with the mediator complex, cell cycle regulators, as well as splicing regulators ^33^. To understand the direct effect of additional understudied mutations on the molecular landscape of these cells, we next compared *Nf1;Mll3* to *Nf1* cells, and *Rb1;Zc3h13* to *Rb1* cells (Figure 7E). We found that 522 genes were upregulated in *Nf1;Mll3* compared to *Nf1* cells, and 175 downregulated (Figure 7E, Table S23). Gene ontology analysis of the gene set associated with higher expression in *Nf1;Mll3* cells revealed significant enrichment of extracellular matrix, EGF-like region, biological adhesion, cell adhesion, calcium ion binding, tube development, growth factor binding, and Wnt receptor signaling pathway (Figure 7F). Comparing *Rb1;Zc3h13* to *Rb1* cells revealed that 703 upregulated and 166 downregulated (Figure 7G, Table S24). *Rb1;Zc3h13*-high genes were enriched in functional categories such as extracellular matrix, immune response, cell adhesion, 2-5-oligoadenylate synthetase, cell morphogenesis, GTPase activity, cell motion, and vasculature development (Figure 7H). Collectively, these findings indicate that the addition of an *Mll3* mutation significantly alters the transcriptome of *Nf1* mutant cells, as does the addition of a *Zc3h13* mutation on *Rb1* mutant cells. Moreover, the differentially expressed genes are associated with pathways that are important for tumorigenesis and/or invasion in the native brain microenvironment (e.g. extracellular matrix, cell adhesion, immune response, vasculature, differentiation, and developmental pathways).

As GBM remains a very challenging cancer type to treat, understanding the molecular changes underlying drug response is of profound importance. We thus performed drug-treatment-RNA-Seq experiments to investigate the transcriptome responses of AAV-CRISPR-induced GBM cells (*Rb1, Rb1;Pten* and *Rb1;Zc3h13*) to TMZ, a chemotherapy agent with significant albeit small survival benefit for GBM patients, among the only 4 currently approved drugs for this disease (Methods) (Figure 8A). Drug response phenotyping showed that *Zc3h13* LOF rendered *Rb1* cells significantly more resistant to TMZ, similar of *Pten* LOF (two tailed t-test, *p* < 0.0001, *Rb1;Pten* vs. *Rb1* and *Rb1;Zc3h13* vs. *Rb1*, for both doses) (Figure 8B-C). Given the differential responses between these three genotypes, we performed mRNA-Seq to profile the transcriptome of these mutant cells treated with TMZ as compared to DMSO controls (Tables S20, S21). Unsupervised hierarchical clustering of the drug-treatment-RNA-Seq data revealed that different biological replicates largely clustered together (Figure S8B), indicating a robust and consistent transcriptional signature associated with each genotype and treatment combination. Comparing TMZ and DMSO treated *Rb1* mutant cells, the expression levels of 352 genes were significantly induced upon TMZ treatment, while 332 genes were reduced (Figure 8D, Table S25). In parallel, TMZ induced 345 while reduced 313 genes in *Rb1;Pten* cells (Figure 8E, Table S26), and induced 703 while reduced 166 genes in *Rb1;Zc3h13* cells (Figure 8F, Table S27). Collectively, the differentially expressed genes in the TMZ vs. DMSO comparisons uncovered a molecular map of the significant transcriptomic differences between genotypes in response to TMZ treatment (Figure 8G).

**Figure 8.**
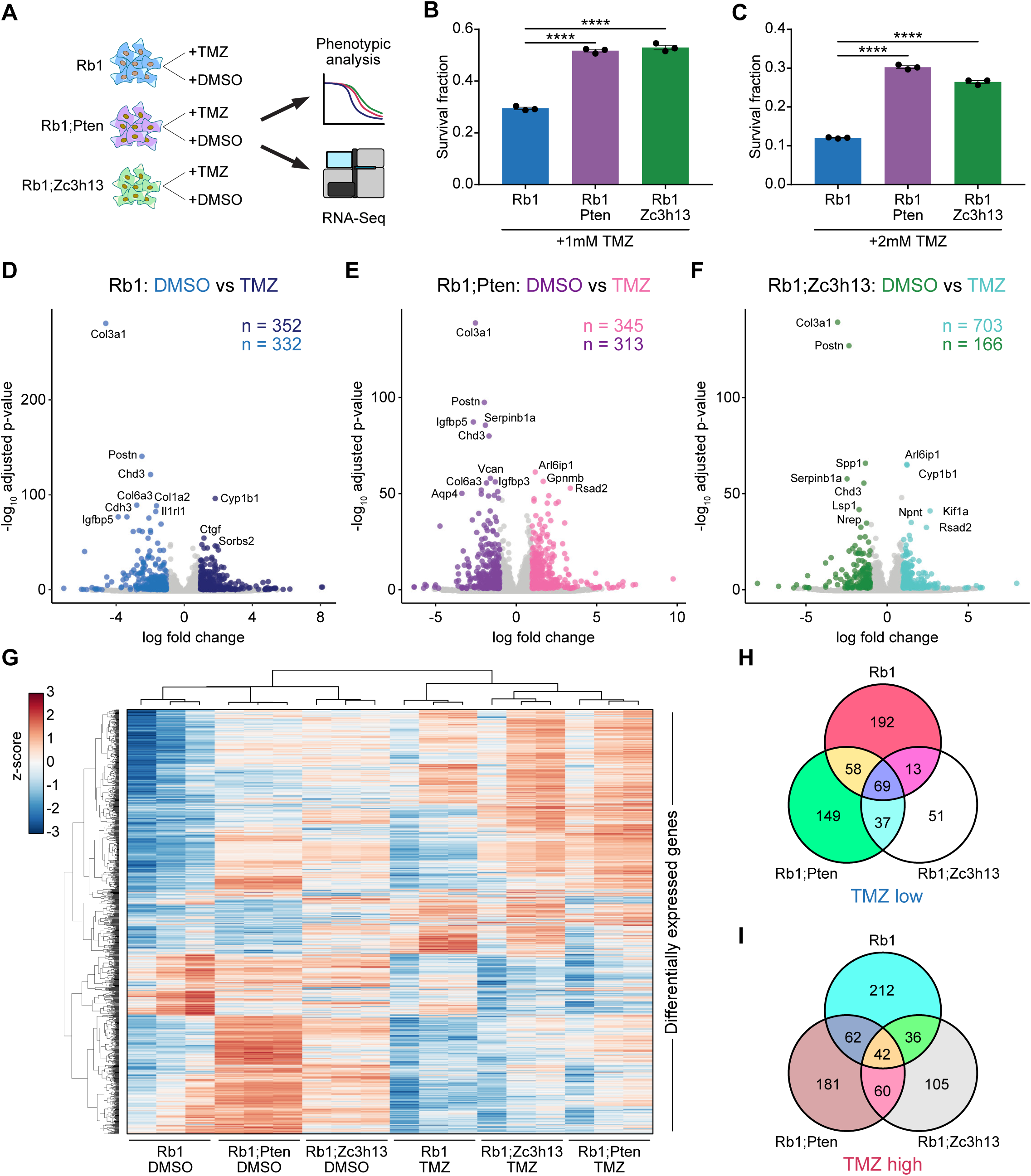
Transcriptome profiling of mouse GBM driver combinations in the presence and absence of a chemotherapeutic agent. **A.** Schematic of GBM-TMZ-RNAseq experimental design. *Rb1, Rb1;Pten,* and *Rb1;Zc3h13* cells were treated with either temozolomide (TMZ) or DMSO. Cells were subjected to phenotypic analysis and RNA-Seq. **B.** Survival fraction of *Rb1* (blue), *Rb1;Pten* (purple), and *Rb1;Zc3h13* (green) cells with 1mM TMZ treatment. *Rb1;Pten* and *Rb1;Zc3h13* cells had significantly higher survival fractions than *Rb1* cells (twosided t-test, **** *p* < 0.0001). **C.** Survival fraction of *Rb1* (blue), *Rb1;Pten* (purple), and *Rb1;Zc3h13* (green) cells with 2mM TMZ treatment. *Rb1;Pten* and *Rb1;Zc3h13* cells had significantly higher survival fractions than *Rb1* cells (twosided t-test, **** *p* < 0.0001). **D.** Volcano plot comparing *Rb1* cells treated with TMZ (dark blue) or DMSO (blue). 352 genes were significantly higher in TMZ-treated cells (Benjamini-Hochberg adjusted *p* < 0.05 and log fold change ≥ 1), and 332 genes were significantly higher in DMSO-treated cells (adjusted *p* < 0.05 and log fold change ≤ −1). Examples of highly differentially expressed genes were labeled. **E.** Volcano plot comparing *Rb1;Pten* cells treated with TMZ (pink) or DMSO (purple). 345 genes were significantly higher in TMZ-treated cells (Benjamini-Hochberg adjusted *p* < 0.05 and log fold change ≥ 1), and 313 genes were significantly higher in DMSO-treated cells (adjusted *p* < 0.05 and log fold change ≤ −1). **F.** Volcano plot comparing *Rb1;Zc3h13* cells treated with TMZ (turquoise) or DMSO (green). 703 genes were significantly higher in TMZ-treated cells (Benjamini-Hochberg adjusted *p* < 0.05 and log fold change ≥ 1), and 166 genes were significantly higher in DMSO-treated cells (adjusted *p* < 0.05 and log fold change ≤ −1). Examples of highly differentially expressed genes were labeled. **G.** Heatmap of all differentially expressed genes among the TMZ vs. DMSO comparisons. Clustering was performed by average linkage using Pearson correlations. Values are shown in terms of z-scores, scaled by each gene. Sample conditions are indicated below. **H.** Venn diagram of genes significantly lower in TMZ-treated vs. DMSO-treated cells for each tested genotype. While 69 genes were similarly downregulated among all 3 genotypes upon TMZ treatment, the differential expression signatures were nevertheless distinct, suggesting differential responses to TMZ treatment in downregulated genes. **I.** Venn diagram of genes significantly higher in TMZ-treated vs. DMSO-treated cells for each tested genotype. While 42 genes were similarly upregulated among all 3 genotypes upon TMZ treatment, the differential expression signatures were nevertheless distinct, suggesting differential responses to TMZ treatment in upregulated genes.

To identify the commonalities and differences in transcriptional responses to TMZ among the three genotypes, we directly compared the differential expressed genes after TMZ treatment in *Rb1*, *Rb1;Pten*, and *Rb1;Zc3h13* mutant cells. Of the genes that were significantly reduced upon TMZ treatment in each group, a total of 69 genes were shared among all three genotypes (Figure 8H, Table S28), indicating that these genes are a common transcriptional response to TMZ. Many of these genes are involved with the extracellular matrix, including *Adamts12, Adamts16, Acan, Cthrc1, Col1a1, Col3a1, Col6a1, Col6a2,* and *Col7a1*. As *Rb1;Zc3h13* and *Rb1;Pten* cells exhibited greater survival fractions with TMZ treatment when compared to *Rb1* cells, we also looked for TMZ-reduced transcripts that were specific to *Rb1;Zc3h13* and *Rb1;Pten* cells. We identified 37 genes that were significantly reduced upon TMZ treatment in *Rb1;Zc3h13* and *Rb1;Pten* cells, but not in *Rb1* cells (Figure 8H, Table S28). These included *Ddit4l* and *Spata18*, both of which are associated with cellular response to DNA damage stimulus, as well as Notch pathway components *Heyl* and *Nrarp*. As for the genes that were significantly induced by TMZ, a total of 42 genes were common in all three genotypes compared to their respective DMSO controls (Figure 8I, Table S29), representing a shared TMZ-induced gene signature. A number of these genes are involved with GTP metabolism and GTPase activity (e.g. *Rragd*, and *Tbc1d8*) and DNA damage (e.g. *Gadd45a*), consistent with the mechanism of action for TMZ. Interestingly, we identified 60 genes that were upregulated upon TMZ treatment in *Rb1;Zc3h13* and *Rb1;Pten* cells, but not in *Rb1* cells. These included *Arl6ip1*, which has been shown to suppress cisplatin-induced apoptosis in cervical cancer cells ^34^, and *Cd274* (also known as *PD-L1*), which is the ligand for the inhibitory receptor *PD-1* that is currently a major focus of investigation in cancer immunotherapy ^35^. Taken together, the transcriptomic analyses provide unbiased molecular signatures underlying the increased TMZ-resistance upon *Zc3h13* or *Pten* mutations in *Rb1*-oncotype glioma cells, suggesting that the precise combinations of mutational drivers present in individual GBMs directly influence therapeutic responses.

## Discussion

GBM was first defined by clinicians based on its presumed glial origin and its highly variable morphology ^36^. As GBM is the most frequent and most aggressive malignant primary brain tumor, it is classified as grade IV by the World Health Organization ^1^. Primary or *de novo* GBM is the most common, and typically progresses rapidly without recognizable symptoms ^2^. Secondary GBM can develop from lower-grade diffuse astrocytoma (grade II) and anaplastic astrocytoma (grade III) ^1^^,2^. GBM patients have poor prognosis and short survival ^37^, though patients with *MGMT* promoter methylation do slightly better ^3^. Certain genetic alterations are known to promote transformation of normal cells in the brain into highly malignant glioma cells ^1,8-10^ particularly those occurring in oncogenes such as *EGFR, IDH1, PIK3CA, ERBB2,* and *PDGFRA,* and tumor suppressor genes *NF1, PTEN, RB1,* and *TP53* ^11-15^. However, to date, there has been no unbiased phenotypic picture of which genetic factors and combinations are necessary or sufficient to drive tumorigenesis in the brain.

To find answers to these questions, it is critical to directly test the hypotheses gleaned from cancer genomics in a controlled experimental setting to find causative genes and to quantitatively measure their phenotypes *in vivo*. Determining whether these alterations are *bona fide* driver mutations requires direct *in vivo* testing, but such testing is generally performed one gene at a time in cell lines or mouse models ^38^. Development and applications of pioneering mouse models of GBM have led to profound progress in our understanding of tumor initiation, stem cell populations, progression and therapeutic responses of GBM ^8,11,12,39-47^, but such studies have been limited to small numbers of genes. With recent developments in genome editing utilizing the CRISPR system ^48-52^, it is now possible to directly mutate oncogenes and tumor suppressor genes in somatic cells for modeling genetic events in various cancer types ^53-63^. However, these methods have only been applied to a small number of genes due to the challenges in generating pool-mutated autochthonous diseases models in the target tissue. While high-throughput CRISPR screens have been successfully demonstrated in cell lines for coding genes ^64,65^ and non-coding elements ^66-68^, as well as transplant tumor models ^69^, direct *in vivo* high-throughput mutational analysis of functional cancer drivers in gliomagenesis has been difficult, due to technological challenges and the nature of biological complexity in the mouse brain. A high-throughput, genome editing method for interrogating a large collection of clinically relevant genes would enable parallelized characterizations of entire cancer landscapes.

We have overcome these challenges by developing a focused sgRNA library with AAVs encoding functional elements and sgRNA pools targeting the top pan-cancer putative TSGs, in combination with conditional LSL-Cas9 transgenic mice. With this platform, we demonstrated high-throughput gene editing in multiplexed autochthonous GBM models in fully immunocompetent mice. Our study provides a massively parallel view of tumor suppressors *in vivo*, revealing the relative selective strength of mutations in these genes when competing in the brain, as well as the driver variations between individuals. As CRISPR targeting might lead to off-target effects at other loci, we performed exome sequencing for a subset of the mTSG brain samples (Methods). This dataset revealed unbiased, exome-wide measurement of other mutations in the coding region (Figure S6E, Tables S30, S31). While some of these other mutations might be caused by off-target effects of CRISPR, it is also possible that part of all of these mutations were induced by the unstable genomes of GBM cells during subsequent evolution after the initial mutagenesis.

Across all genes tested, the mutation frequencies in this highly multiplexed genetic mouse model of GBM significantly correlate with their mutation frequencies in human patients from two large independent cohorts (TCGA and Yale Glioma bank), suggesting the clinical relevance of the findings. Several of the novel SMGs highly enriched in this mouse study have also been associated with GBM in the clinical setting with human samples, such as *B2M, CIC, MLL2, MLL3, SETD2, ZC3H13* and *ARID1B* ^15,70-73^. We subsequently validated several of these drivers, alone or in combination, using a minipool AAV-CRISPR approach. To further explore the molecular mechanisms underlying the selection forces that favored these combinations, we investigated several of the gene combinations by transcriptomic analysis, finding hundreds of differentially expressed genes between the two distrinct *Nf1*- and *Rb1*- oncotypes, as well as second mutations (*Mll3, Pten, Zc3h13*) on top of an *Nf1* or *Rb1* mutant background. Moreover, we found that *Zc3h13* mutation significantly enhanced resistance to TMZ, to a similar degree as an additional *Pten* mutation, in the *Rb1* background. Despite being frequently mutated across several human cancers, the function of *Zc3h13* has not been explicitly characterized in the context of cancer. Finally, to better understand the mechanisms by which different driver combinations influence resistance to chemotherapeutic agents such as TMZ, we performed drug-treatment-RNA-Seq and identified multiple common and unique pathways that were induced or reduced upon TMZ treatment. Because differences in driver mutations can dramatically affect treatment efficacies in pre-clinical animal models and in patients ^74,75^, a functional understanding of cancer drivers is therefore essential for precision medicine ^76^.

Compared to lentiviruses, AAVs have their own advantages and limitations. Of note, we also performed a lentiCRISPR direct *in vivo* screen in GBM, using the same mTSG library (Methods, Table S32, Figure S8C-D). Using current standard protocols, we found that the AAV-mTSG CRISPR library resulted in more robust gliomagenesis *in vivo* compared to lentiCRISPR mTSG, in terms latency (death of first animal, 84 vs. 200 days), survival (median survival, 4 vs. 10 months), and penetrance (100% vs. 67%). AAVs usually do not integrate into the genome, except under certain circumstances (low rate of integration at *AAVS1* locus) ^77,78^. Thus, AAV-encoded transgenes such as exogenously supplied sgRNAs do not replicate as cells divide during tumor progression, limiting the readout of the mTSG library by PCR amplification of the sgRNA cassette itself. To sidestep this issue, we achieved successful readout of driver mutations in the targeted genes by directly sequencing the predicted sgRNA cutting sites for indels using ultra-deep targeted captured sequencing. Although this approach is currently limited in the number of sgRNAs that can be read out given the number of cells transduced, the amount of genomic DNA needed, and the nature of capture sequencing technology, we anticipate this method can be further improved by increasing the power of customized capture probes and sequencing at even greater depths. Thus, our approach of AAV-CRISPR mediated pooled mutagenesis in conjunction with targeted capture sequencing provides an efficient platform for massively paralleled analysis of GBM drivers directly *in vivo*.

One of the key advantages of this approach is the ability to perform high-throughput analysis of mutants in an autochthonous model of GBM; that is, models in which the tumors directly evolve from normal cells at the organ site *in situ* in immunocompetent mice and without cellular transplantation. This platform can be readily extended to study other types of cancer for tumor progression, as well as therapeutic responses *in vivo*. While the present study was focused on tumor suppressors in GBM, the AAV-CRISPR approach can be readily adapted for the study of oncogenes. The use of catalytically-dead Cas9 ^79-81^, catalytically dead gRNAs ^82^, CRISPR-targeted cytidine deaminases ^83^, or point mutation via HDR ^54^, in conjunction with the pooled AAV-CRISPR platform presented in this study, will open up the potential for oncogene screens, either through overexpression/amplification using CRISPRa or by inducing precise point mutations usingor HDR or CRISPR-targeted cytidine deaminases. Taken together, our study provides a systematic and unbiased molecular landscape of functional tumor suppressors in an autochthonous mouse model of GBM, opening new paths for high-throughput analysis of cancer gene phenotypes directly *in vivo*.

## Acknowledgments

We thank all members in Chen, Sharp, Zhang and Platt laboratories, as well as various colleagues in Yale Department of Genetics, Systems Biology Institute, Yale Cancer Center and Stem Cell Center, Koch Institute and Broad Institute at MIT for assistance and/or discussion. We thank the Center for Genome Analysis, Center for Molecular Discovery, High Performance Computing Center, West Campus Analytical Chemistry Core and West Campus Imaging Core and Keck Biotechnology Resource Laboratory at Yale, as well as Swanson Biotechnology Center at MIT, for technical support.

SC is supported by Yale University Systems Biology Institute Startup Fund, Damon Runyon Research Foundation Dale Frey Award for Breakthrough Scientists (DRG-2117-12; DFS-13-15), Melanoma Research Foundation (MRA-412806), St-Baldrick’s Foundation, Yale Institutional Research Grant from American Cancer Society, Yale Skin Cancer SPORE, Yale Lung Cancer SPORE, Breast Cancer Alliance, and NIH/NCI Center for Cancer Systems Biology (U54). RJP is supported by National Center of Competence in Research Molecular Systems Engineering (NCCRMSE) and ETH Zurich; and partially by the McGovern Institute and NSF (1122374). PAS is supported by NIH (R01-CA133404) and the Lambert Fund. F.Z. is supported by the NIH / NIMH (5DP1-MH100706 and 1R01-MH110049), NSF, NY Stem Cell Foundation, HHMI, Poitras, Simons, Paul G. Allen Family, and Vallee Foundations, D.R. Cheng and B. Metcalfe. F.Z. is an NY Stem Cell Foundation-Robertson Investigator. CDG, PR are supported by the NIH Graduate Training Grant (T32GM007499). RDC, MBD and MWY are supported by the NIH MSTP training grant (T32GM007205). F.S. is supported by NCCRMSE and ETH Zurich. GW is supported by RJ Anderson Postdoctoral Fellowship.

## Contributions

SC and RJP conceived, designed the study and performed the initial set of experiments. CDG, GW, RJP and SC performed the majority of animal work and histology. RDC developed the algorithms and performed integrative analyses of all the data. GW performed exome-capture, mutant cell line generation, drug treatment and RNA-Seq. FS performed AAV production. SC performed MRI. MWY contributed to data analysis. LY, YE, MBD, MAM, SZ and PR contributed to experiments including mouse breeding, genotyping, cloning, cell culture, virus prep, injection, necropsy and sample prep. KB assisted captured and exome sequencing. MG provided clinical insights. PAS, FZ, RJP, and SC jointly supervised the work. RDC and SC wrote the manuscript with inputs from all authors.

## Methods

### Design, synthesis and cloning of the mTSG library

Due to cellular complexity in the organs *in vivo*, we set out to design a small focused library for viral transduction. Briefly, pan-cancer mutation data from 15 cancer types were retrieved from The Cancer Genome Atlas (TCGA portal) via cBioPortal ^84^ and Synapse (https://www.synapse.org). Significantly mutated genes were calculated similar to previously described methods ^28-31^. Known oncogenes were excluded and only known or predicted tumor suppressor genes (TSGs) were included. The top 50 TSGs were chosen, and their mouse homologs (mTSG) were retrieved from mouse genome informatics (MGI) (http://www.informatics.jax.org). A total of 49 mTSGs were found. A total of 7 known housekeeping genes were chosen as internal controls. We designed sgRNAs against these 56 genes using a previously described method ^64,65^ with our custom scripts. Five sgRNAs were chosen for each gene, plus 8 non-targeting controls (NTCs), making a total 288 sgRNAs in the mTSG library. There were two sets of duplicate sgRNAs, Cdkn2a-sg2 / Cdkn2a-sg5, and Rpl22-sg4 / Rpl22-sg5, leaving a total of 286 unique sgRNAs.

### Design, cloning of an AAV-CRISPR GBM vector and mTSG sgRNA library cloning

An AAV-CRISPR vector was designed for astrocyte-specific genome editing. This vector contains a cassette specifically expresses Cre recombinase under the control of a *GFAP* promoter for conditional induction of Cas9 expression in astrocytes in the brain when delivered to LSL-Cas9 mice ^54^. Two sgRNA cassettes were built in this vector, one encoding an sgRNA targeting *Trp53*, the most frequently mutated gene in cancer ^28,29,31^, with the other being an empty sgRNA cassette (double SapI sites for sgRNA cloning) enabling flexible targeting of genes of interest in either individual or pooled manner. The vector was generated by gBlock gene fragment synthesis (IDT) followed by Gibson assembly (NEB). The mTSG libraries were generated by oligo synthesis, pooled, and cloned into the double SapI sites of the AAV-CRISPR GBM vector. The library cloning was done at over 100x coverage to ensure proper representation. Plasmid library representation was readout by barcoded Illumina sequencing as described previously ^69^ with primers customized to this vector.

### AAV-mTSG viral library production

The AAV-CRISPR GBM plasmid vector (AAV-vector) and library (AAV-mTSG) were subjected to AAV9 production and chemical purification. Briefly, HEK 293FT cells (ThermoFisher) were transiently transfected with transfer (AAV-vector or AAV-mTSG), serotype (AAV9) and packaging (pDF6) plasmids using polyethyleneimine (PEI). Each replicate consist of five of 80% confluent HEK 293FT cells in 15-cm tissue culture dishes or T-175 flasks (Corning). Multiple replicates were pooled to enhance production yield. Approximately 72 hours post transfection, cells were dislodged and transferred to a conical tube in sterile PBS. 1/10 volume of pure chloroform was added and the mixture was incubated at 37°C and vigorously shaken for 1 hour. NaCl was added to a final concentration of 1 M and the mixture was shaken until dissolved and then pelleted at 20k g at 4°C for 15 minutes. The chloroform layer was discarded while the aqueous layer was transferred to another tube. PEG8000 was added to 10% (w/v) and shaken until dissolved. The mixture was incubated at 4°C for 1 hour and then spun at 20k g at 4° C for 15 minutes. The supernatant was discarded and the pellet was resuspended in DPBS plus MgCl_2_ and treated with Benzonase (Sigma) and incubated at 37°C for 30 minutes. Chloroform (1:1 volume) was then added, shaken, and spun down at 12k g at 4C for 15 min. The aqueous layer was isolated and passed through a 100 kDa MWCO (Millipore). The concentrated solution was washed with PBS and the filtration process was repeated. Virus was titered by qPCR using custom Taqman assays (ThermoFisher) targeted to Cre.

### Design, cloning of lentiCRISPR GBM vectors and mTSG sgRNA library, and lentivirus production

Two lentiCRISPR vectors were designed, one for constitutive, and the other for astrocyte-specific genome editing. These vectors contain a cassette specifically expresses Cre recombinase under the control of an *EFS* promoter or a *GFAP* promoter for conditional induction of Cas9 expression in the brain when deliver to LSL-Cas9 mice. Two sgRNA cassettes were built in this vector, one encoding an sgRNA targeting *Trp53*, with the other being an empty sgRNA cassette (double BsmbI sites for sgRNA cloning) enabling flexible targeting of genes of interest in either individual or pooled manner. These vectors were generated by gBlock gene fragment synthesis (IDT) followed by Gibson assembly (NEB). The mTSG libraries were generated by oligo synthesis, pooled, and cloned into the double BsmbI sites of the lentiCRISPR GBM vectors. The library cloning was done at over 100x coverage to ensure proper representation. Plasmid library representation was readout by barcoded Illumina sequencing as described above, with primers customized to the vectors. The LentiCRISPR GBM plasmid vector (Lenti-vector) and library (Lenti-mTSG) were subjected to high-titre lentivirus production and purification. Briefly, HEK 293FT cells (ThermoFisher) were transiently transfected with transfer (Lenti-vector or Lenti-mTSG), and packaging (psPAX and pMD2.G) plasmids using PEI or Lipofectamine. Each replicate consist of five of 80% confluent HEK 293FT cells in 15-cm tissue culture dishes or T-175 flasks (Corning). Multiple replicates were pooled to enhance production yield. Approximately 48 hours post transfection, virus-containing media was collected, and purified via sucrose gradient ultracentrifugation at >=30,000 rpm for 2-3h. The supernatant was discarded and the pellet was dried and resuspended with 100 μl sterile PBS in 4 °C overnight. Virus was titered by viral protein p24 ELISA (RnD).

### Animal work statements

All animal work was performed under the guidelines of Yale University Institutional Animal Care and Use Committee (IACUC) and Massachusetts Institute of Technology Committee for Animal Care (CAC), with approved protocols (Chen-2015-20068, Zhang-0414-024-17, and Sharp-0914-091-17), and were consistent with the Guide for Care and Use of Laboratory Animals, National Research Council, 1996 (institutional animal welfare assurance no. A-3125-01).

### Stereotaxic surgery and virus transduction in the mouse brain

Conditional LSL-Cas9 knock-in mice were breed in a mixed 129/C57BL/6 background. Mixed gender (randomized males and females) 6-14 week old mice were used in the experiment. Animals were maintained and breed in standard individualized cages with maximum of 5 mice per cage, with regular room temperature (65-75°F, or 18-23°C), 40-60% humidity, and a 12h:12h light cycle. Mice were anesthetized by intraperitoneally injection of ketamine (100 mg/kg) and xylazine (10 mg/kg), or by inhalation of isoflurane at approximately 2% for 20-30 minutes. We also administered buprenorphine HCl (0.1 mg/kg), or carprofen (5.0 mg/kg) intraperitoneally as a pre-emptive analgesic. Reflexes were tested before surgical procedures. Once subject mice were in deep anesthesia, they were immobilized in a stereotaxic apparatus (Kopf, or Stoelting) using intra-aural positioning studs and a tooth bar to immobilize the skull. Heat is provided for warmth by a standard heating pad, or a heatlamp. According to the mouse brain stereotaxic coordinates ^85^, we drilled a 1-2 mm hole on the surface of the skull, and used a 33 G Nanofil syringe needle (World Precision Instrument) to inject into the lateral ventricle (LV) at 0.6-1.0 mm caudal/posterior to Bregma, 0.8-1.5 mm right-side lateral to Bregma, and 2.0-3.0 mm deep from the pial surface for injection (coordinates: A/P −0.6 to −1.0, M/L 0.8 to 1.5, D/V −2.0 to −3.0). For a small fraction of animals, injections were made into hippocampus (HPF) at the following coordinates (A/P −1.3, M/L 0.6, D/V −1.7). PBS, 8 uL AAV (Between 1 x 10^10^ – 1x 10^11^ viral genome copies, or Cre copy number equivalent), or 8 uL lentivirus (Between 8 x 10^9^ – 8 x 10^10^ viral particles, or p24 equivalent) was injected into the right hemisphere of the brain for each mouse. Injection rates were monitored by an UltraMicroPump3 (World Precision Instruments). After injection, the incision site was closed with 6-0 Ethilon sutures (Ethicon by Johnson & Johnson), or a VetBond tissue glue (3M). Animal were postoperatively hydrated with 1 mL lactated Ringer’s solution (subcutaneous) and housed on warmed cages or in a temperature controlled (37°C) environment until achieving ambulatory recovery. Meloxicam (1-2 mg/kg) was also administered subcutaneously directly after surgery.

### MRI

MRI imaging was performed using standard imaging protocol with MRI machines (Varian 7T/310/ASR-whole mouse MRI system, or Bruker 9.4T horizontal small animal systems). Briefly, animals were anesthetized using isoflurane, and setup in the imaging bed with a nosecone providing constant isoflurane. A total of 20 − 30 views were acquired for each mouse brain using a custom setting: echo time (TE) = 20, repetition time (TR) = 2000, slicing = 0.5 mm. Raw image stacks were processed using Osirix or Slicer tools ^86^. Rendering and quantification were performed using Slicer (www.slicer.org). For all mice with brain tumors, only 1 tumor was observed per mouse. Tumors were approximate as spheres and their sizes were calculated with the following formula:

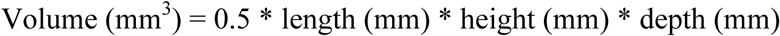

### Survival analysis

We observed that mice that developed brain tumors rapidly deteriorate in their body condition scores. Mice with observed macrocephaly and body condition score ≤ 1 were euthanized and the euthanasia date was recorded as the last survival date. Occasionally mice bearing brain tumors died unexpectedly early, and the date of death was recorded as the last survival date. Cohorts of mice stereotaxically injected with PBS, AAV-vector or AAV-mTSG virus were monitored for their survival. Survival analysis was analyzed using standard Kaplan – Meier method. Of note, several AAV-vector or PBS injected mice were sacrificed at time points earlier than 299 days (at times when a certain AAV-mTSG mice were found dead or euthanized due to poor body conditions), to provide time-matched histology, but those mice are healthy without brain tumor or other signs of detectable symptoms. Mice euthanized early in healthy state were excluded from calculation of survival percentage.

### Mouse brain dissection, fluorescent imaging, and histology

Mice were sacrificed by carbon dioxide asphyxiation or deep anesthesia with isoflurane followed by cervical dislocation. Mouse brains were manually dissected under a fluorescent stereoscope (Zeiss, Olympus or Leica). Brightfield and/or GFP fluorescent images were taken for the dissected brain, and overlaid using ImageJ ^87^. Brains were then fixed in 4% formaldehyde or 10% formalin for 48 to 96 hours, embedded in paraffin, sectioned at 6 μm and stained with hematoxylin and eosin (H&E) for pathology. For tumor size quantification, H&E slides were scanned using an Aperio digital slidescanner (Leica). Tumors were manually outlined as region-of-interest (ROI), and subsequently quantified using ImageScope (Leica). Sections were de-waxed, rehydrated and stained using standard immunohistochemistry (IHC) protocols as previously ^69,88^. The following antibodies were used for IHC: rabbit anti-Ki67 (abcam ab16667, 1:500), rabbit anti-GFAP (Dako, 1:500), and mouse anti-Cas9 (Diagenode, 1:300).

### Mouse tissue collection for molecular biology

Mouse brain (targeting organ) and liver (non-targeting organ) were dissected and collected manually. For molecular biology, tissues were flash frozen with liquid nitrogen, ground in 24 Well Polyethylene Vials with metal beads in a GenoGrinder machine (OPS diagnostics). Homogenized tissues were used for DNA/RNA/protein extractions using standard molecular biology protocols.

### Genomic DNA extraction from cells and mouse tissues

For genomic DNA (gDNA) extraction, 50-200 mg of frozen ground tissue was resuspended in 6 ml of Lysis Buffer (50 mM Tris, 50 mM EDTA, 1% SDS, pH 8) in a 15 ml conical tube, and 30 μl of 20 mg/ml Proteinase K (Qiagen) was added to the tissue/cell sample and incubated at 55 °C overnight. The next day, 30 μl of 10 mg/ml RNAse A (Qiagen) was added to the lysed sample, which was then inverted 25 times and incubated at 37 °C for 30 minutes. Samples were cooled on ice before addition of 2 ml of pre-chilled 7.5M ammonium acetate (Sigma) to precipitate proteins. The samples were vortexed at high speed for 20 seconds and then centrifuged at ≥ 4,000 x *g* for 10 minutes. Then, a tight pellet was visible in each tube and the supernatant was carefully decanted into a new 15 ml conical tube. Then 6 ml 100% isopropanol was added to the tube, inverted 50 times and centrifuged at ≥4,000 x *g* for 10 minutes. Genomic DNA was visible as a small white pellet in each tube. The supernatant was discarded, 6 ml of freshly prepared 70% ethanol was added, the tube was inverted 10 times, and then centrifuged at ≥4,000 x *g* for 1 minute. The supernatant was discarded by pouring; the tube was briefly spun, and remaining ethanol was removed using a P200 pipette. After air-drying for 10-30 minutes, the DNA changed appearance from a milky white pellet to slightly translucent. Then, 500 μl of ddH_2_O was added, the tube was incubated at 65 °C for 1 hour and at room temperature overnight to fully resuspend the DNA. The next day, the gDNA samples were vortexed briefly. The gDNA concentration was measured using a Nanodrop (Thermo Scientific).

### Targeted capture sequencing probe design

Targeted capture sequencing probes were designed as following: the predicted cutting sites (3bp 5’ of PAM) of the 280 gene-targeting sgRNAs in the mTSG library plus the *Trp53*-targeting sgRNA in the vector were retrieved from mouse genome (mm10). The 140bp sequences of the flanking regions of the cutting sites (5’- 70bp and 3’- 70bp) were retrieved using Bedtools ^89^. The regions were consolidated using NimbleDesign (Roche / NimbleGen), and probe matches were set with these parameters (Preferred Close Matches = 3, where initial selection of probes for a given region will only include probes with 3 close matches or less; and Maximum Close Matches = 20, where if there are insufficient probes available for a given region at the Preferred Close Match number, the threshold will be incrementally increased to 20 until adequate coverage is achieved. After consolidation, a number of 178 regions covering 277 sgRNAs, with a total of 33638bp were covered in the probeset, with Target Bases Covered = 32239 (95.8%) and one target sgRNA without coverage due to a lack of qualified candidate probes in the region.

### Targeted capture sequencing

The mTSG-Amplicon targeted capture sequencing probes were synthesized using the SeqCap EZ Probe Pool synthesis procedure (Roche). The capture sequencing was done following standard Illumina-Roche-Illumina protocols. Genomic DNA samples from mouse organs were subjected to fragmentation, followed by a library preparation step using KAPA Library Preparation Kit (Illumina). The libraries were then amplified using LM-PCR, hybridized to the mTSG-Amplicon probe pool, washed and recovered, and amplified with multiplexing barcodes using LM-PCR. The multiplexed library was then QC’ed using qPCR, and subjected to high-throughput sequencing using the Hiseq-2500 or Hiseq-4000 platforms (Illumina) at Yale Center for Genome Analysis. 277/278 (99.6%) of unique targeted sgRNAs were captured for all samples from this experiment, with the missing one being Arid1a-sg5.

### mTSG sgRNA cutting efficiency measurement

The mouse mTSG sgRNA cutting efficiency measurement was performed similar to the screen with the exception of early sampling. Briefly, AAV-mTSG library virus was injected in to LV of LSL-Cas9 mice, but instead of end-point tumor, mice were sacrificed at early time point (3.5 weeks post injection) and examined under fluorescent stereoscope, and GFP+ regions from the brain were dissected. Genomic DNA was extracted, and subjected to capture sequencing.

### Mouse whole-exome capture sequencing

The mouse whole-exome capture was performed using SeqCap EZ exome kit (Roche). Briefly, capture sequencing was done following standard Illumina-Roche-Illumina protocols. Genomic DNA samples from mouse organs were subjected to fragmentation, followed by a library preparation step using KAPA Library Preparation Kit (Illumina). The libraries were then amplified using LM-PCR, hybridized to the exome probe pool, washed and recovered, and amplified with multiplexing barcodes using LM-PCR. The multiplexed library was then QC’ed using qPCR, and subjected to high-throughput sequencing using the Hiseq-2500 or Hiseq-4000 platforms (Illumina) at Yale Center for Genome Analysis.

### Illumina sequencing data processing and variant calling

FASTQ reads were mapped to the mm10 genome using the bwa mem function in BWA v0.7.13 ^90^. Bam files were merged, sorted, and indexed using bamtools v2.4.0 ^91^ and samtools v1.3 ^92^. For each sample, indel variants were called using samtools and VarScan v2.3.9 ^93^. Specifically, we used samtools mpileup (–d 1000000000 –B –q 10), and piped the output to VarScan pileup2indel (–min-coverage 1 –min-reads2 1 –min-var-freq 0.001 –p-value 0.05). To link each indel to the sgRNA that most likely caused the mutation, we took the center position of each indel and mapped it to the closest sgRNA cut site.

### Calling significantly mutated sgRNAs and significantly mutated genes

We further filtered all detected indels by requiring that each indel must overlap the ± 3 base pairs flanking the closest sgRNA cut site, as Cas9-induced double-strand breaks are expected to occur within a narrow window of the predicted cut site. We then utilized a series of criteria to identify high-confidence mutations: 1) As an initial pass to exclude possible germline mutations, we removed any sgRNAs with indels present in more than half of the control samples with greater than 5% variant frequency. In our data, this filter specifically removed Rps19_sg5 from further consideration. 2) To determine significantly cutting sgRNAs in each sample, we used a false-discovery approach based on the PBS and vector control samples. For each sgRNA, we first took the highest % variant read frequency across all control samples; in order for a mutation to be called in an mTSG sample, the % variant read frequency had to exceed the control sample cutoff. However, since the base vector contained a *Trp53* sgRNA (*Trp53* sg8) whose cut site was only 1 bp away from the target site of *Trp53* sg4 (from mTSG library), we only considered PBS samples when calculating the false-discovery cutoff for *Trp53* sg4. Nevertheless, in the current study this exception was unnecessary because of our third filter: 3) As we were most interested in identifying the dominant clones in each sample, we further set a 5% variant frequency cutoff on top of the false-discovery cutoff. These criteria gave us a binary table (i.e. not significantly mutated vs. significantly mutated) detailing each sgRNA and whether its target site was significantly mutated in each sample. None of the AAV-vector samples passed the 5% cutoff at the p53 sg4/8 target site, which is consistent with our observation that no tumors were found in vector-treated animals. To convert significantly cutting sgRNAs into significantly mutated genes, we simply collapsed the binary sgRNA scores by gene, such that if any of the sgRNAs for a gene were found to be significantly cutting, the gene would be called as significantly mutated.

### Exome sequencing data analysis

For exome sequencing analysis, we imposed a modified set of criteria on each detected variant: 1) ≥ 10 supporting reads for the reference allele; 2) ≥ 10 supporting reads for the variant allele; 3) the variant is within ± 6 bp of a Cas9 PAM, NGG (or CCN on the reverse strand); 4) a variant allele frequency < 75%, as this was the maximum detected variant frequency out of the mTSG brain samples; 5) the variant was not detected in any sequenced control samples, which were considered as germline variants.

### Clustering of variant frequencies to infer clonality of tumors

For each mTSG brain sample, we extracted the individual variants that comprised the SMS calls in that sample, with a cutoff of 5% variant frequency to eliminate low-abundance variants. Because of these cutoffs, 3 sequenced mTSG brain samples were not eligible for variant frequency clustering analysis. To identify clusters of variant frequencies in an unbiased manner, we modeled the variant frequency distribution with a Gaussian kernel density estimate, using the Sheather-Jones method to select the smoothing bandwidth. From the kernel density estimate, we then identified the number of local maxima (i.e. “peaks”) within the density function. The number of peaks thus represents the number of variant frequency clusters for an individual sample, which is an approximation for the clonality of the tumors.

### Coding frame analysis

For coding frame and exonic/intronic analysis, we only considered the indels that were associated with a sgRNA which had been considered significantly mutated in that particular sample. This final set of significant indels was converted to .avinput format and subsequently annotated using ANNOVAR v. 2016Feb01, using default settings ^94^.

### Co-occurrence and correlation analysis

Co-occurrence analysis was performed by first generating a double-mutant count table for each pairwise combination of genes in the mTSG library. Statistical significance of the co-occurrence was assessed by hypergeometric test. For correlation analysis, we first collapsed the % variant frequency tables on the gene level (in other words, summing the % variant frequencies for all 5 of the targeting sgRNAs for each gene). Using these summed % variant frequency values, we calculated the Spearman correlation between all gene pairs, across each mTSG sample. Statistical significance of the correlation was determined by converting the correlation coefficient to a t-statistic, and then using the t-distribution to find the associated probability. A similar approach was used to analyze co-occurring mutations in human TCGA GBM data.

### Testing driver combinations with sgRNA minipool

Mixtures of five sgRNAs targeting each gene were cloned as sgRNA minipool into the same astrocyte-specific AAV-CRISPR vector. For gene pair targeting, the five-sgRNA single gene minipools from both genes were mixed 1:1. Plasmid mixes were then packaged into AAV1/2. Briefly, HEK293FT cells were transfected with the minipools plasmids, pAAV1 plasmid, pAAV2 plasmid, helper plasmid pDF6, and PEI Max (Polysciences, Inc. 24765-2) in DMEM (ThermoFisher, 10569-010). 72 hours post transfection, cell culture media was discarded and cells were rinsed and pelleted via low speed centrifugation. Cells were then lysed and the supernatant containing viruses was applied to HiTrap heparin columns (GE Biosciences 17-0406-01) and washed with a series of salt solutions with increasing molarities. During the final stages, the eluates from the heparin columns were concentrated using Amicon ultra-15 centrifugal filter units (Millipore UFC910024). Titering of viral particles was executed by quantitative PCR using custom Cre-targeted Taqman probes (ThermoFisher). After packaging, AAV minipools were stereotaxically injecting into the ventricle of LSL-Cas9 mice. Survival and histology analysis followed injection as described above. Several control (uninjected, EYFP and vector) mice were sacrifice as surrogate histology although they were in good body condition and were subsequently found devoid of tumor.

### Generation of Nf1 and Rb1 mutant cell lines from primary GBMs induced by AAV-CRISPR minipools

Autochthonous mouse GBMs were induced by stereotaxic injection with the *Nf1* or *Rb1* AAV minipool (in the AAV9-sg*Trp53*-sgX-GFAP-Cre vector described above). Tumor-containing brains were visually inspected under a fluorescent dissecting scope, made into single-cell suspension through physical dissociation plus Collagenase/DNase digestion, and cultured in DMEM supplemented with 10% FBS and Pen/Strep. Growing clones were further established as autochthonous mouse GBM cell lines.

### Single sgRNA knockout lentiviral production

Lenti-pHKO-U6-sgBsmBI-EF1a-Puro-P2A-FLuc was generated by subcloning P2A-Fluc expression cassette into lentiviral CRISPR knockout vector by Gibson assembly. For the cloning of sgRNA targeting individual genes such as *Pten*, *Arid1b*, *Mll3 (Kmt2c)*, *B2m*, and *Zc3h13* (Table S1), the corresponding oligos were synthesized, annealed and cloned into BsmBI linearized lentiviral knockout vector. Lentiviruses were produced by transfecting lentiviral knockout plasmids, together with pMD2.G and psPAX2 into 80-90% confluent HEK293FT cells, with viral supernatants collected 48 and 72 h posttransfection, aliquoted and stored in −80°C.

### Generation of NF1+geneX and RB1+geneX knockout cell lines

The *Nf1* and *Rb1* knockout tumor cells were infected by single sgRNA knockout lentiviruses at M.O.I <= 0.3 to further knockout desired geneX. 24 h post-infection, lentiviral transduced cells were selected by the addition of 4-8 μg/ml puromycin and were split 2-3 days.

### Temozolomide (TMZ) treatment, Cell viability assay, and RNAseq

After 7-9 days’ culture under puromycin selection, lentiviruses-infected *Nf1* and *Rb1* knockout tumor cells were plated in triplicates into 96-well plate at a density of 2.5 x 10^3^ cells per well, and ~5 x 10^6^ cells were collected at the same time and used for cutting efficiency analysis. One day after plating, either TMZ or DMSO was added at a concentration of 10 μM, 100 μM, 500 μM, 1 mM, and 2 mM. After 3 days of drug/vehicle treatment, cell viability was measured using CellTiter Glo (Promega) according to the manufactures’ protocol. Briefly, we first equilibrated the CellTiter Glo at room temperature for 1 h before use. The media of 96-well plates was aspirated, and then 50 μL fresh DMEM+10%FBS and 50 μL CellTiter Glo was added. The luminescent signals were readout using EnVision plate reader (PerkinElmer). For RNAseq samples preparation, cell lines harboring specific gene knockouts were cultured for 7-9 days under the selection pressure of puromycin, and then plated into 6-well plates at a density of 2 x 10^5^ cells per well in triplicates. 24 h after plating, 1 mM TMZ or DMSO in fresh DEME + 10% FBS was added and cultured for another 48 h. Then, cellular RNA of control or treated cells were extracted by adding 350 μL TRIzol Reagent (Invitrogen) directly into 6-well plates to lyse the cells, followed by gently shaking the plates and incubation for 5-10 min to complete and lysis and homogenization. Then, 70 μL chloroform was added, vigorously mixed, and centrifuged at 16,000 g for 15 min. Transfer the RNA containing aqueous phase to a new tube and further purified using RNeasy mini kit (Qiagen). After eluting RNA from column using Nuclease free water, the concentrations of sample RNA were normalized into 150-300 ng/uL for RNAseq.

### T7 Endonuclease I (T7E1) Assays

The genomic DNA of these cells that were collected after 9 days of puromycin selection was extracted by using QuickExtract™ DNA Extraction Solution (Epicentre), mixed well and incubated at 65°C for 30-60 min. Then, 1-2 μL of genomic DNA from parental or Lenti-sgRNA transduced cells was used as the template to amplify gene of interest using surveyor primers (Table S33) with thermocycling conditions as 98°C for 1 min, 35 cycles of (98°C for 1 s, 60°C for 5 s, 72°C for 10 s), and 72°C for 1 min. The PCR products were gel-purified using QIAquick Gel Extraction Kit from 2% E-gel EX and quantified, followed by PCR products denaturing at 95°C for 5 min, and annealing by using following conditions: ramp from 95 to 85 °C at a rate of −2°C per seconds; from 85°C to 25°C at a rate of −0.1°C per second, and 4°C hold. 1 μL of T7 endonuclease I was added into annealed oligo, and incubated at 37°C for 60 min to digest the mismatched sites. The digested PCR products were loaded into 2% E-gel EX, and the amount of DNA fragments were quantified. The cutting efficiency was calculated to estimate gene editing using the following formula: Indels (%) = 100 x (1 – (1- fraction cleaved)1/2).

### Transcriptome profiling of different driver combinations in the presence and absence of chemotherapy

Mixtures of five sgRNAs targeting each gene were cloned as sgRNA minipool into the same astrocyte-specific AAV-CRISPR vector. After packaging, AAV minipools were stereotaxically injecting into the lateral ventricle of LSL-Cas9 mice. Cell lines were derived from mouse GBMs by single-cell isolation, plating and culture in DMEM media. Additional driver mutations were introduced by lentiCRISPR where applicable. GBM cells with different drivers were treated with DMSO or TMZ for 48h, and harvested for mRNA-seq for transcriptome profiling. Briefly, total RNA was extracted from cancer cells derived from AAV-CRISPR minipools induced GBM treated with DMSO or TMZ, using commercially available kits (Qiagen / Thermofisher). PolyA-mRNA library was constructed using Illumina TruSeq mRNA library prep kit, and sequenced on Illumina Hiseq 2500 and/or Hiseq 4000 platform.

### RNA-seq differential expression analysis

Strand-specific single-end RNA-seq read files were analyzed to obtain transcript level counts using Kallisto ^95^, with the settings –rf-stranded -b 100 and subsequently used the *tximport* R package to collapse to gene level counts. Pairwise differential expression analysis between groups was then performed using edgeR with default settings ^96^.

### Pathway enrichment analysis of differentially expressed transcripts

Using an adjusted p-value cutoff of 0.05, and a log fold change threshold of ±1, we determined the set of genes that were significantly upregulated or downregulated. We then used the resultant gene sets for DAVID functional annotation analysis ^97^. We considered a GO category as statistically significant if the Benjamini-Hochberg adjusted p-value was less than 0.05.

### GBM comparative cancer genomics analysis using TCGA datasets

Somatic mutation calls, copy number variation calls, RNA-seq expression z-scores, and clinical data containing patient survival information were obtained through cBioPortal for GBM on November 15, 2016. Pearson correlation coefficients were calculated comparing mouse and human mutation frequencies; statistical significance was calculated by converting the correlation coefficient to a t-statistic, and then using the t-distribution to calculate significance.

### GBM comparative cancer genomics analysis using Yale Glioma datasets

Somatic mutation calls and copy number variation calls and partial clinical data containing diagnostic information were obtained from Yale Glioma tissue bank and data bank. All patient samples were deidentified. The general description, demographics and tumor characteristics were in supplemental tables. Total event for each patient was calculated as the sum of mutation events and copy number variant events. Pearson correlation coefficients were calculated comparing mouse and human mutation frequencies; statistical significance was calculated by converting the correlation coefficient to a t-statistic, and then using the t-distribution to calculate significance.

### Histology analysis of clinical GBM samples from Yale Glioma tissue bank

Histology sections were obtained from Yale Glioma tissue bank. All patient samples were de-identified. The mutations associated with specific samples were obtained from Yale Glioma data bank. Slides stained with H&E or anti-GFAP were subsequently scanned using a slidescanner (Leica) and subjected to pathological analysis.

### Blinding statement

Investigators were blinded for histology scoring, captured sequencing and RNA-seq, but not blinded for dissection, MRI or survival analysis.

### Code availability

Custom scripts used to process and analyze the data will be deposited to public code-sharing domain (GibHub) and are available upon request.

### Accession

Genomic sequencing data are in the process of being deposited in NCBI SRA under a pending accession number. CRISPR reagents (AAV-CRISPR and lentiCRISPR backbone plasmids and mTSG libraries) are available to the academic community through Addgene.

## Figure legends

**Figure S1.**
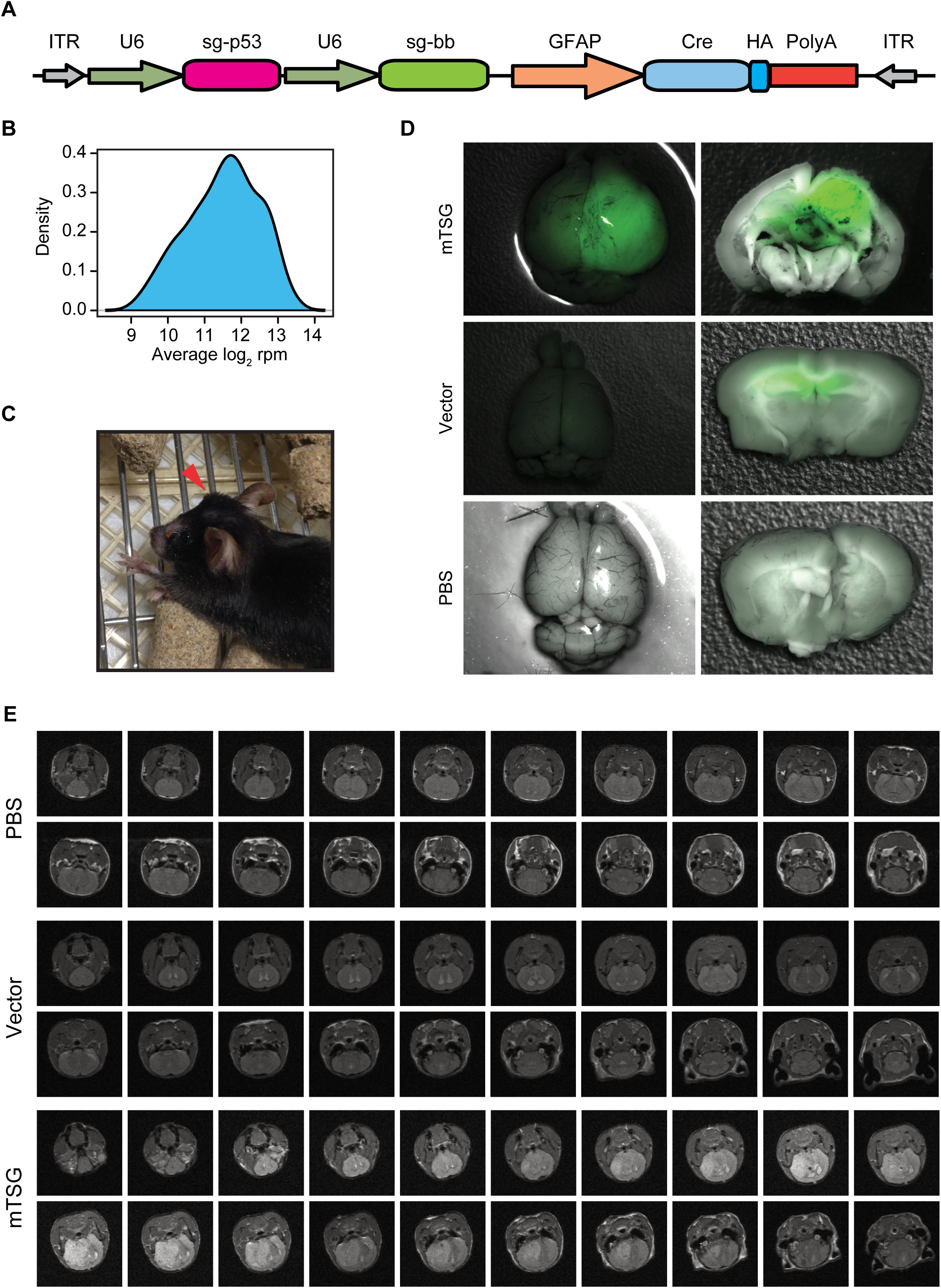
Additional data of massively parallel GBM suppressor analysis by AAV-CRISPR library mediated pooled mutagenesis. **A.** Schematic of the AAV vector used in the study. The vector contains a cassette expressing Cre recombinase under a GFAP promoter, a *p53* sgRNA under U6 promoter, and an empty cassette for expression of custom cloned sgRNA(s). **B.** Plasmid library representation of the AAV-CRISPR mTSG library. **C.** A representative AAV-mTSG injected mouse showing macrocephaly. **D.** Dissected whole brains from PBS, AAV-vector and AAV-mTSG injected mice (left) and sections (right) visualized under a fluorescent stereoscope. **E**. Full-spectrum MRI series of representative mouse brains in PBS, vector and mTSG group. Mice under anesthesia were imaged with a small animal MRI imaging system. 20 MRI sections are shown for each condition. Brain tumors were found in AAV-mTSG injected mice but not in matched PBS or AAV-vector injected mice.

**Figure S2.**
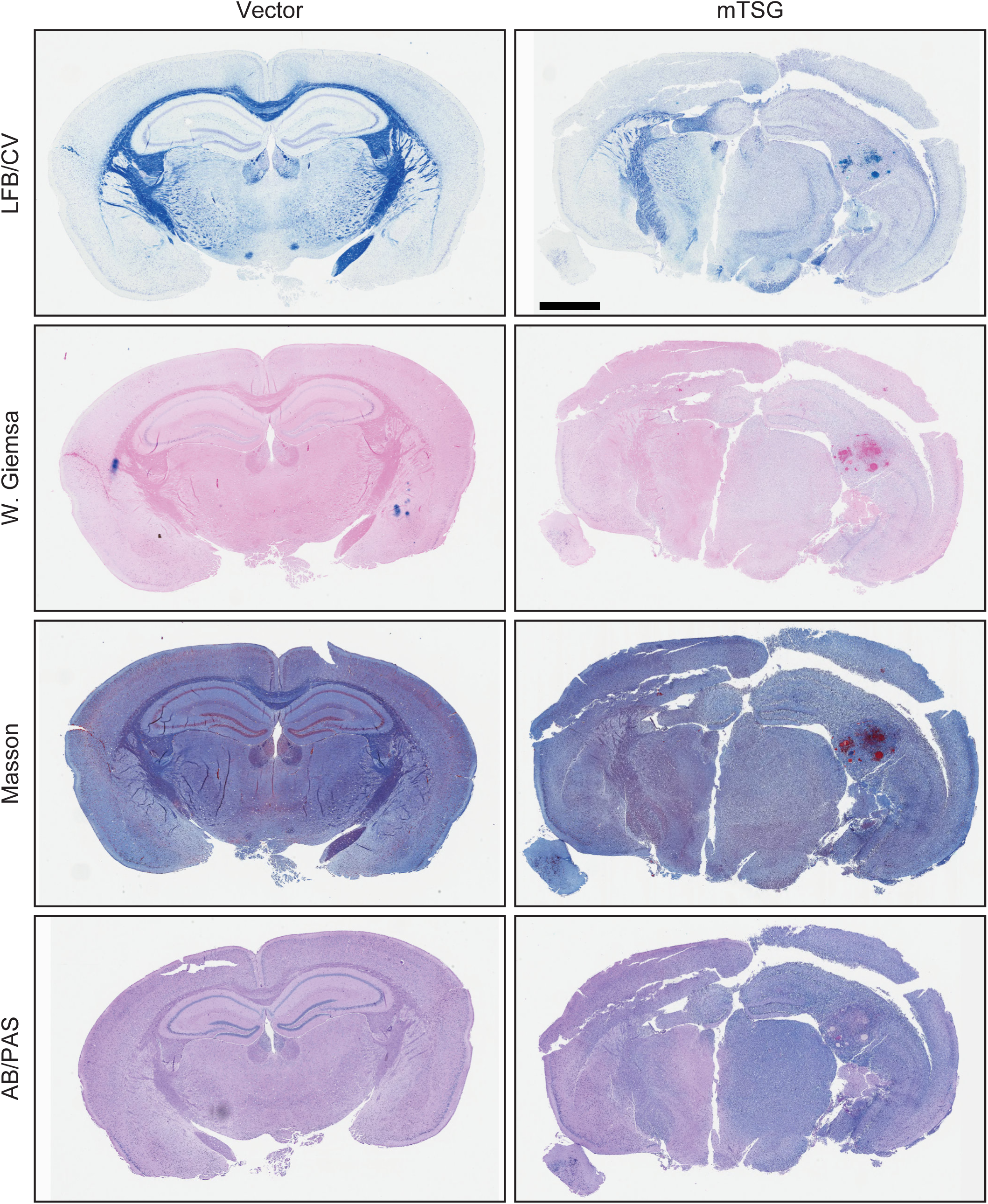
Full-scan histology images of special staining of mouse brain sections in vector and mTSG groups. Panels from top to bottom: Luxol fast blue Cresyl violet (LFB/CV) staining, Wight Giemsa staining, Masson staining and Alcian blue Periodic acid – Schiff (AB/PAS) staining of representative mouse brain sections in vector and mTSG groups. Scale bar = 1 mm.

**Figure S3.**
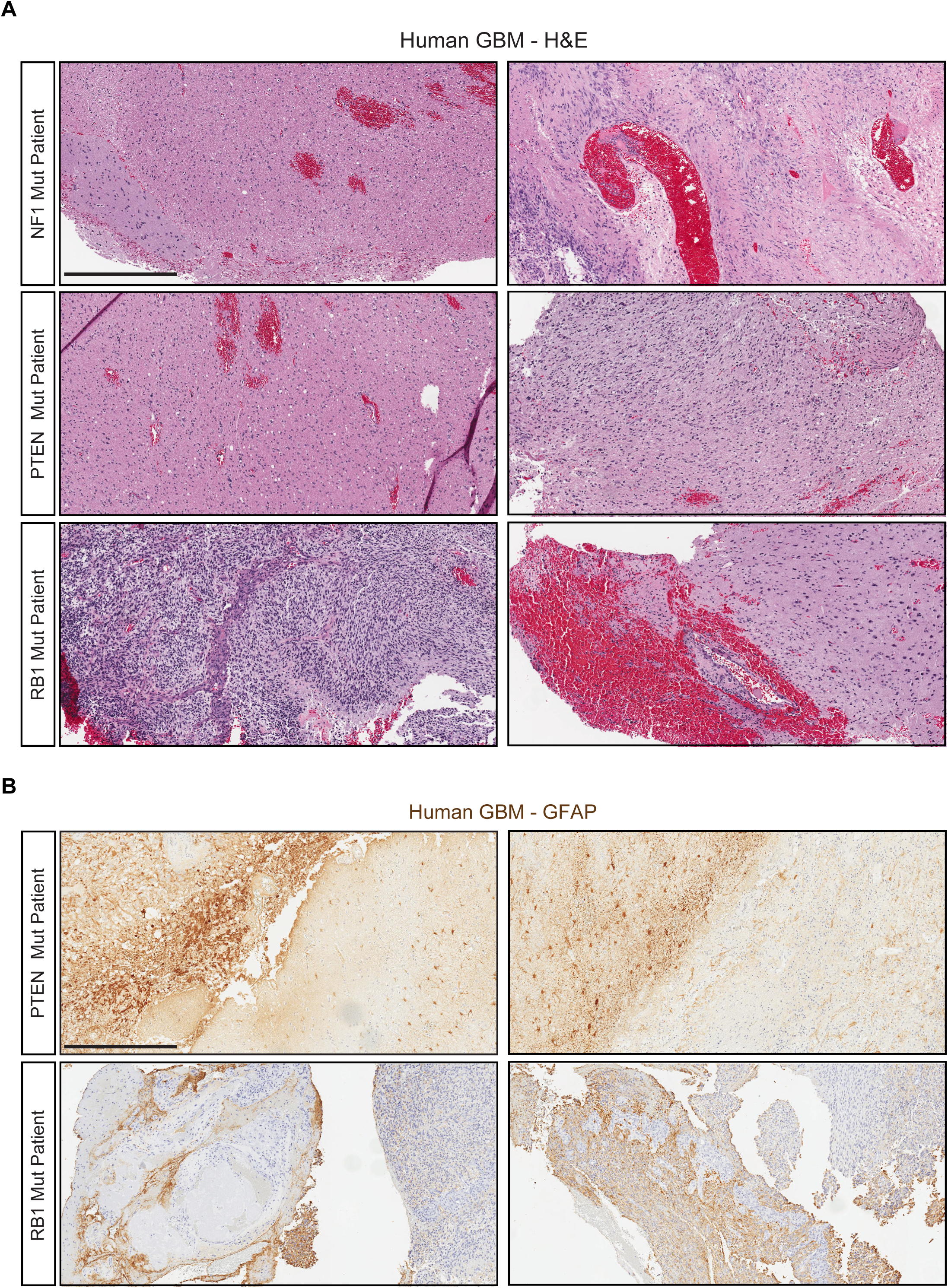
Representative histopathology images of human GBM. **A.** Representative images of H&E stained brain sections from human GBM patient samples from Yale Glioma tissue bank. Images from the three rows represent GBM with significant mutations in *NF1, PTEN* and *RB1,* respectively. Pathological features such as giant aneuploid cells with pleomorphic nuclei, angiogenesis, necrosis and hemorrhage were evident in these tumors. Scale bar = 0.5 mm. **B.** Representative images of anti-GFAP stained brain sections from human GBM patient samples from Yale Glioma tissue bank. Images from the two rows represent GBM with significant mutations in *PTEN* and *RB1,* respectively. *PTEN* tumors were mostly GFAP-positive. *RB1* tumors have mixtures of GFAP-positive and GFAP-negative cells. *NF1* tumors were not shown due to availability of GFAP staining sections. Scale bar = 0.5 mm.

**Figure S4.**
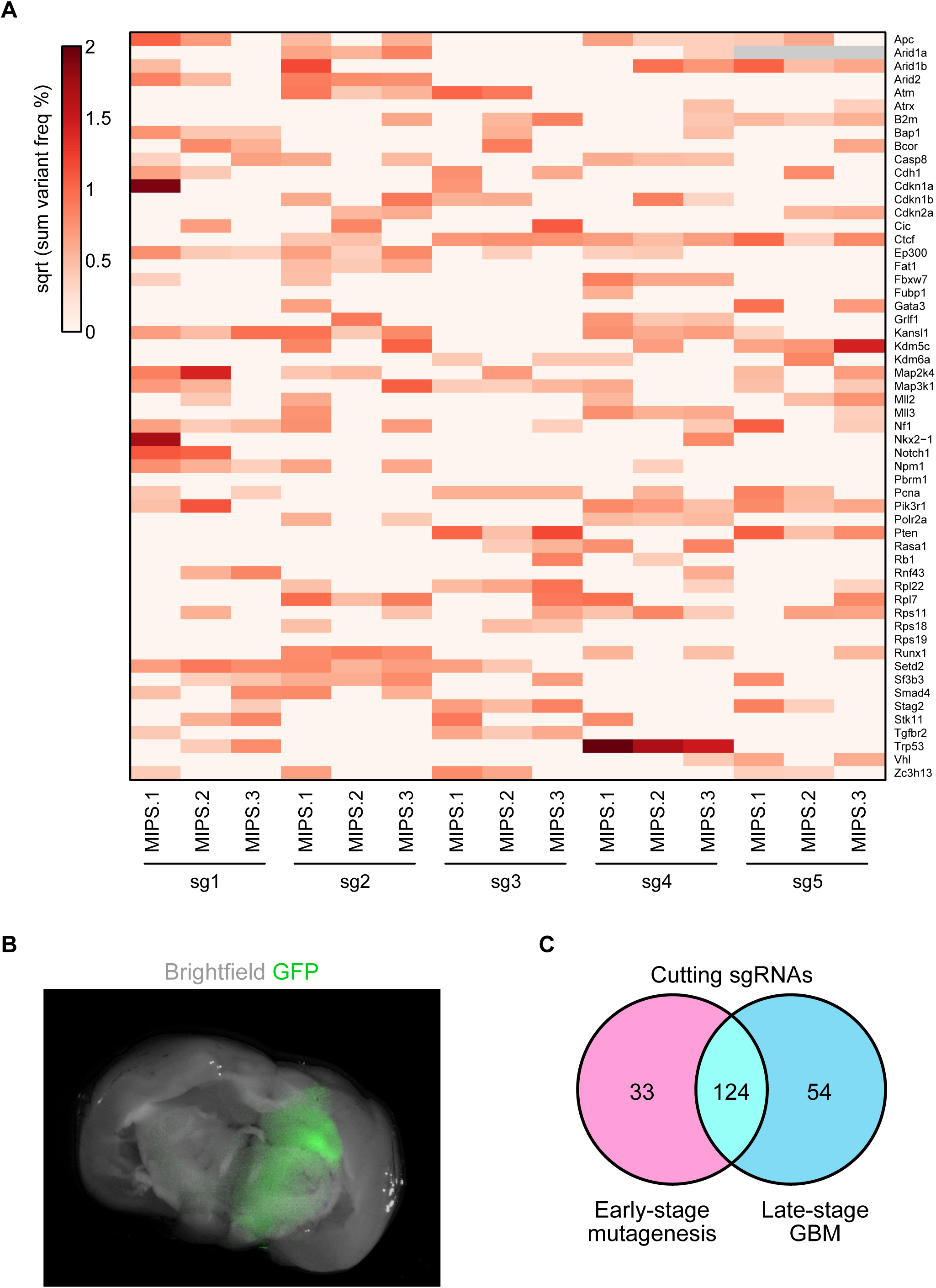
Early time point analysis of sgRNA cutting efficiency by molecular inversion probe sequencing. **A.** Heatmap of sum variant frequencies for each sgRNA across the 3 *in vivo* infection replicates. Each row denotes one gene, while each column corresponds to a specific sgRNA and replicate. Variant frequencies are square-rooted to improve visibility. **B.** Dissected whole brain from an AAV-mTSG injected mouse for early time point analysis, visualized under a fluorescent stereoscope. GFP (green) is shown as an overlay on the brightfield image. **C.** Venn diagram detailing the overlap between cutting sgRNAs identified in early-stage mutagenesis and late-stage GBMs. Differences in the identified cutting sgRNAs were due to differential selection pressures, insufficient time for CRISPR mutagenesis to occur in early time point brains, and/or allele frequencies below detection limit of capture sequencing.

**Figure S5.**
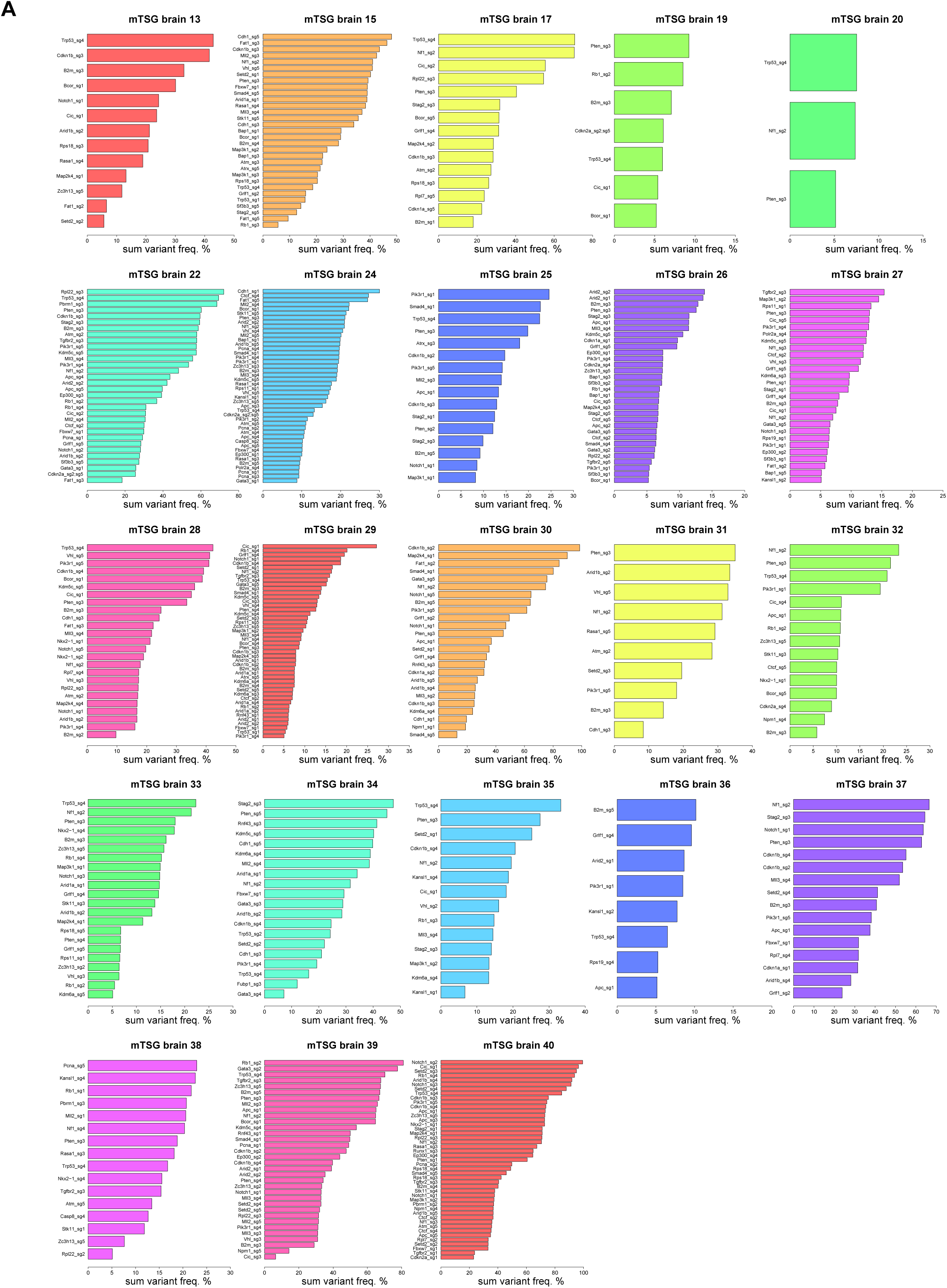
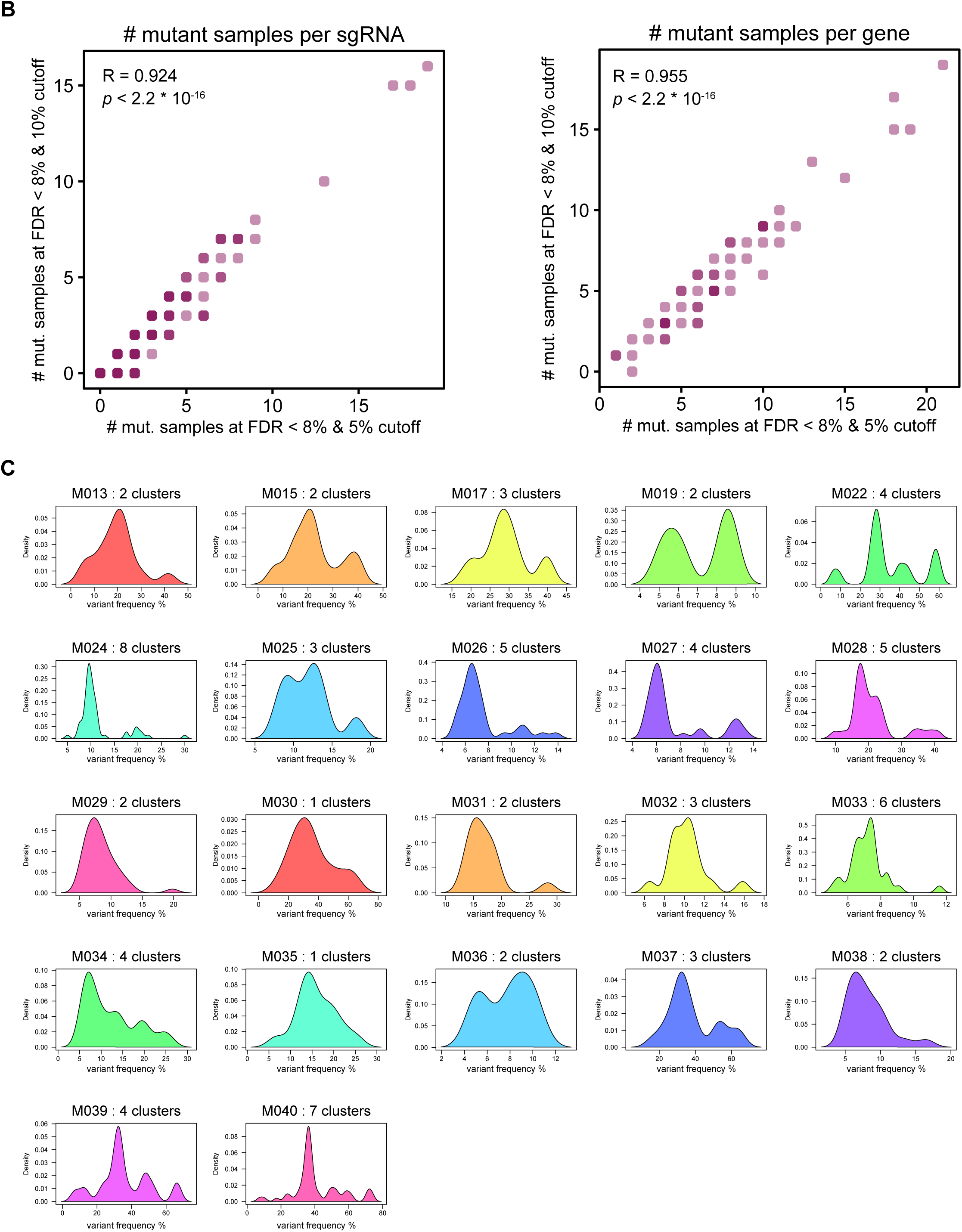
Mutational oncotypes of all GBM mice induced with AAV-CRISPR mTSG library. **A.** Waterfall plots of significantly mutated sgRNA sites across all mTSG brain samples, sorted by sum variant frequency. Two samples (mTSG brain 11, mTSG brain 21) are not shown, as these samples were not found to have any significantly mutated sgRNA sites per our stringent variant calling strategy. The extensive mutational landscape in these samples shows strong positive selection for LOF in gliomagenesis in the brains of these mice. **B.** Scatterplots of the number of samples with an SMS call per sgRNA (left) or SMG call per gene (right), using two different thresholds for calling SMSs. In conjunction with the FDR approach, the use of either a flat 5% or 10% variant frequency cutoff did not affect the results at either the sgRNA or gene level. The Spearman correlation coefficients and associated p-values are shown on the plot. **C.** Gaussian kernel density estimate of variant frequencies within each mTSG brain sample. The number of peaks in the kernel density estimate is an approximation for the clonality of each sample. From this analysis, most (20/22) samples appeared to be composed of multiple clones, with only two (M030, M035) monoclonal samples. Of note, 3/25 sequenced mTSG brain samples did not have sufficient highfrequency variants for clustering analysis.

**Figure S6.**
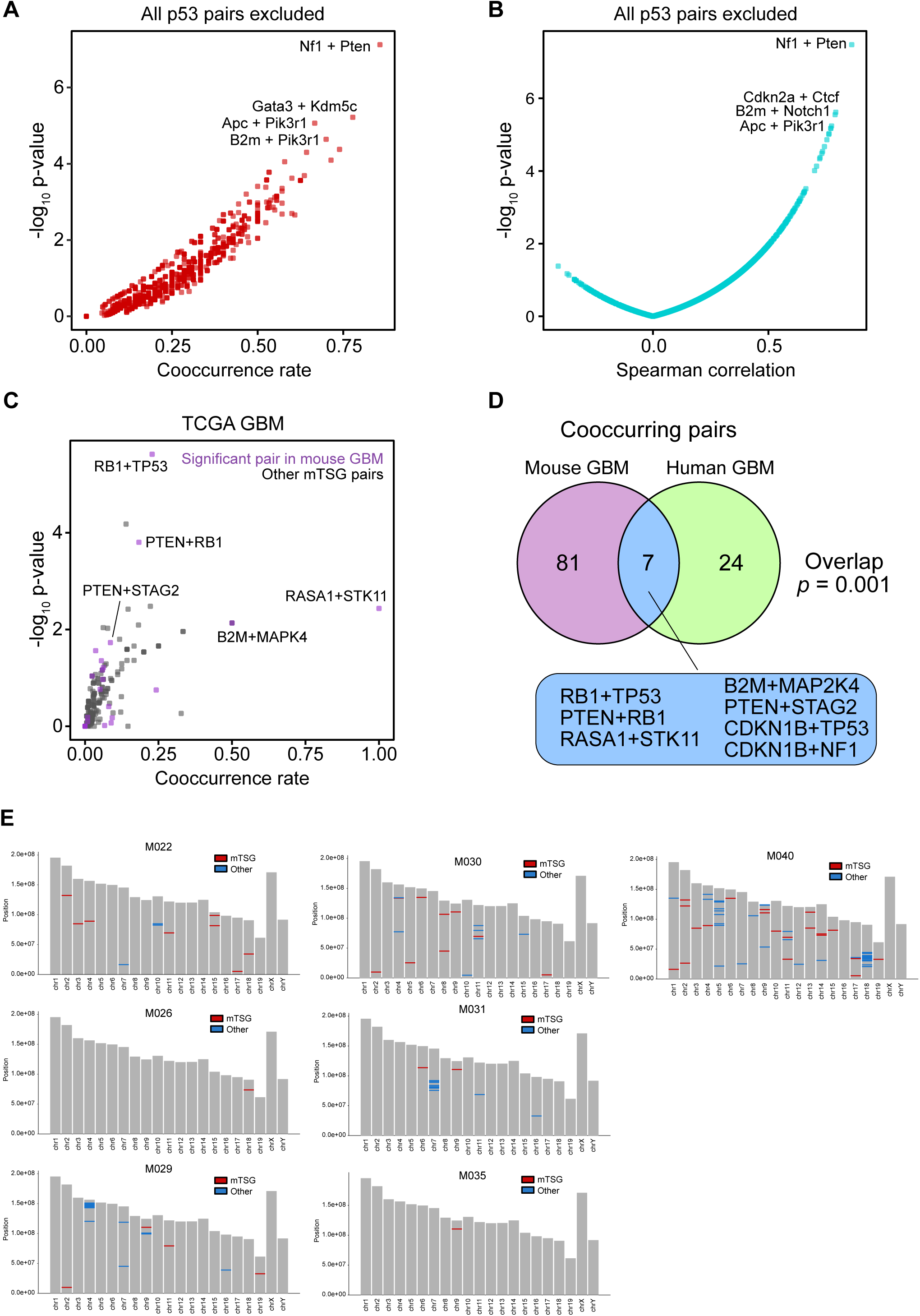
Additional analysis of co-mutated pairs. **A.** Scatterplot of the cooccurrence rate of a given mutation pair, plotted against −log_10_ p-values. Representative co-occurring pairs are indicated. All pairs involving *Trp53* were excluded from this analysis. **B.** Scatterplot of pairwise Spearman correlations plotted against −log_10_ p-values. Representative top pairs are indicated. All pairs involving *Trp53* were excluded from this analysis. **C.** Scatterplot of the cooccurrence rate of a given mutation pair in the TCGA human GBM dataset, plotted against −log_10_ p-values. Top co-occurring pairs are indicated. **D.** Venn diagram of co-occurring pairs identified in mouse GBM (Benjamini-Hochberg adjusted *p* < 0.05, either cooccurrence or Spearman correlation analysis) and/or in human GBM (*p* < 0.05). 7 gene pairs were found to be significant in both mouse and human GBM. The overlap between the two datasets was significant (hypergeometric test, *p* = 0.001). **E.** Whole-exome analysis of possible off-target mutations generated by AAV-CRISPR mTSG. Chromosomal map of potential off-targets in AAV-CRISPR mTSG brain samples. Indels in mTSG genes are marked in red, while possible off-target mutations are marked in blue.

**Figure S7.**
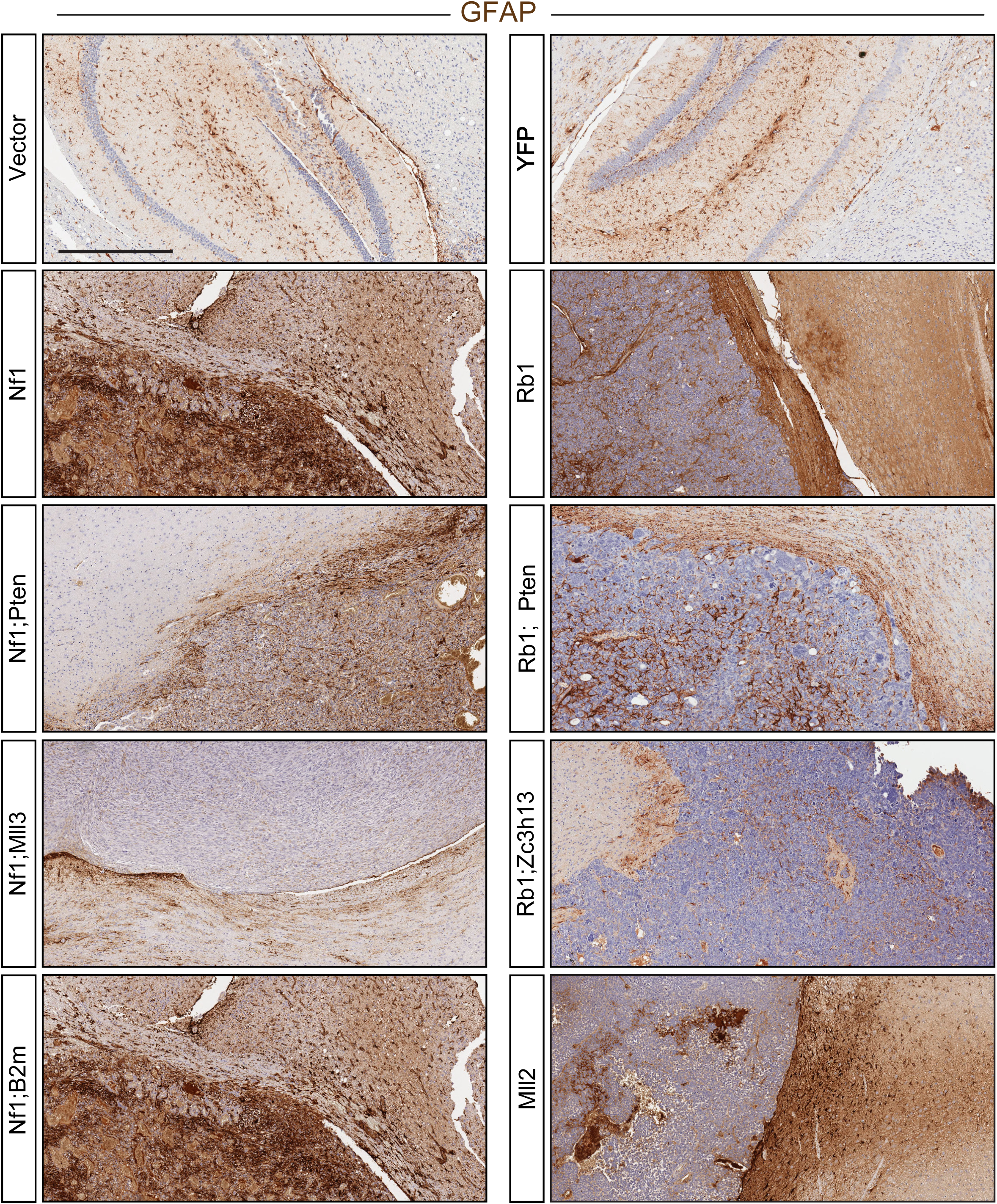
GFAP immunohistochemical characterization of brain sections from mice treated with AAV sgRNA minipools. GFAP immunohistochemistry of brain sections from mice treated with various AVV minipools. Brain tumors in *Nf1, Nf1;Pten,* and *Nf1;B2m* mice were strongly positive for GFAP, while tumors in *Nf1;Mll3* mice were positive at an intermediate level. Brain tumors in *Rb1, Rb1;Pten,* and *Rb1;Zc3h13* mice contained a mixture of GFAP positive and negative cells, similar to the GFAP staining pattern with human patient GBM samples. Brain tumors in *Mll2* mice were variably GFAP positive. Scale bar = 0.5 mm.

**Figure S8.**
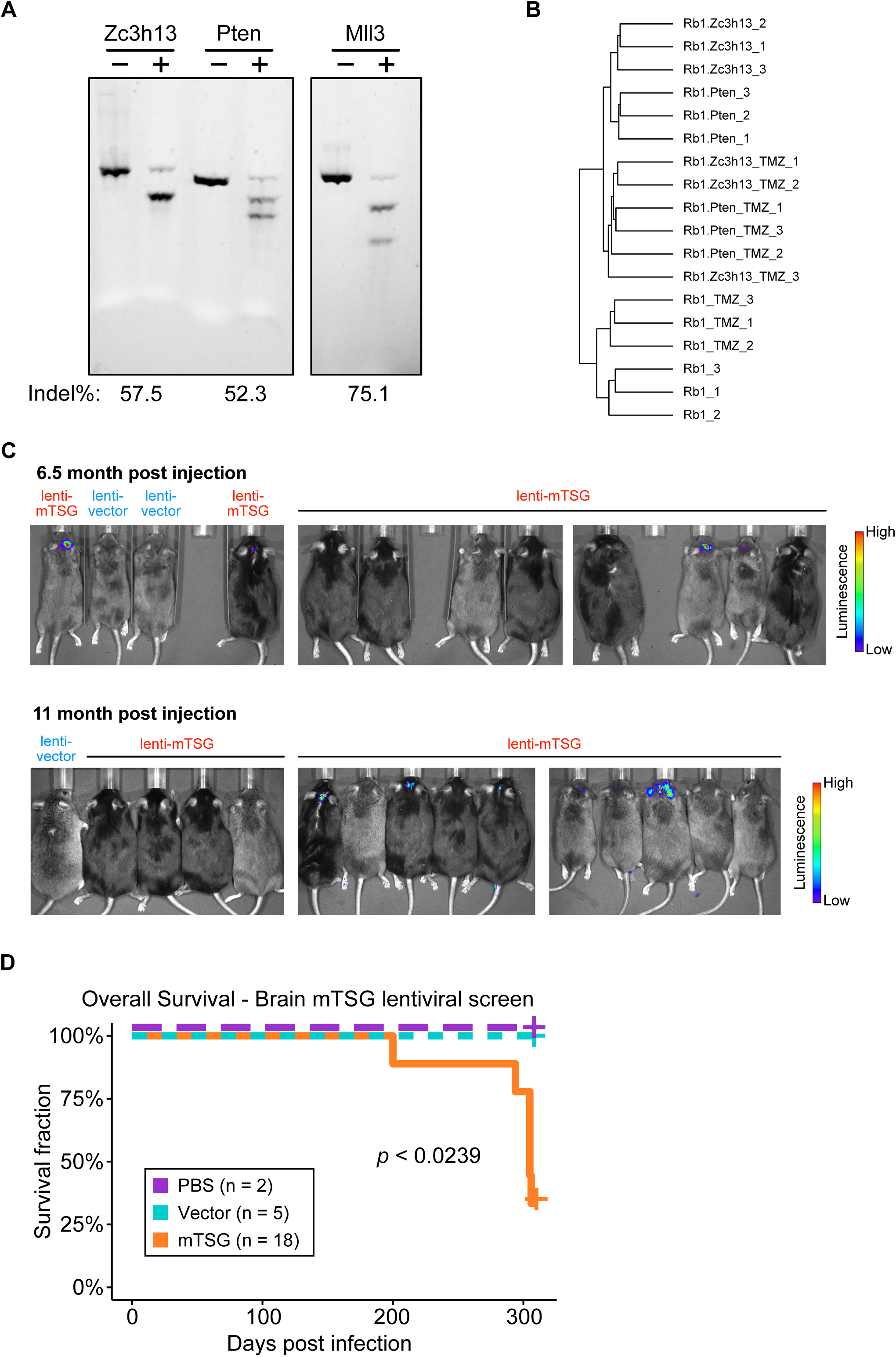
Additional supplemental data related to the study. **(A-B) Additional data for RNA-Seq experiments** **A.** T7E1 nuclease assay to confirm mutagenesis by CRISPR/Cas9 at the indicated target gene. The indel frequency is indicated at the bottom of the gel image. **B.** Hierarchical clustering of *Rb1, Rb1;Pten,* and *Rb1;Zc3h13* cell samples treated with DMSO or TMZ. Biological replicates of the same genotype and treatment condition largely clustered together, indicating high reproducibility between biological replicates. **(C-D). LentiCRISPR mTSG direct in vivo GBM screen** **C.** IVIS imaging of mice injected with lenti-vector or lenti-mTSG library, showing luminescence in the brains of a fraction of lenti-mTSG injected mice, but not in vector injected mice. Mice were imaged at 6.5 months post injection (mpi), where 4/18 mice imaged were luciferase positive (10 were shown). These 4 mice were sacrificed as they developed poor body conditions and brain tumors, before the end of 10 mpi. Mice were imaged again at 11 mpi, where 6/14 mice imaged luciferase positive, which were subsequently sacrificed as they developed poor body conditions and brain tumors. **D.** Kaplan-Meier curves for overall survival (OS) of mice injected with PBS (n = 2), lenti-vector (n = 5) or lenti-mTSG library (n = 18). OS for PBS and vector groups were both 100%, where the curves are dashed and slightly offset for visibility. Log-rank (LR) test, *p* < 0.0239, mTSG vs. vector or PBS; LR test, *p* = 1, vector vs. PBS.

## Supplemental Tables

Table S1. SgRNA spacer sequences in the mTSG library

Table S2. MRI tumor size statistics of GBM mice induced with AAV-mTSG library

Table S3. Survival statistics of GBM mice induced with AAV-mTSG library

Table S4. Histology tumor size statistics of GBM mice induced with AAV-mTSG library

Table S5. mTSG-Amplicon capture probe design of targeted region coordinates and coverage

Table S6. Sample metadata for targeted capture sequencing of GBM mice induced with AAV-mTSG library

Table S7. Targeted capture sequencing coverage statistics across all predicted cutting sites of sgRNAs in AAV-mTSG library

Table S8. Raw variant calling of all samples with targeted capture sequencing

Table S9. SgRNA level sum indel frequency table for early time point brain samples as an approximation for sgRNA cutting efficiency

Table S10. SgRNA level sum indel frequency table for all samples with targeted capture sequencing

Table S11. SgRNA level SMS calls in GBM mice induced with AAV-mTSG library

Table S12. Gene level mSMG calls in GBM mice induced with AAV-mTSG mTSG library

Table S13. Comparative mutation frequencies in the GBMs between AAV-mTSG mice and human patients from TCGA GBM cohort

Table S14. De-identified general description, demographics and tumor characteristics of a Glioma patient cohort in Yale Glioma tissue bank.

Table S15. Mutational matrix of all mTSG orthologs in human GBM samples from the database of Yale Glioma tissue bank

Table S16. Comparative mutation frequencies in the GBMs between AAV-mTSG mice and human patients from Yale Glioma cohort

Table S17. Co-occurrence analysis of mSMG pairs in GBM mice induced with AAV-mTSG library

Table S18. Spearman correlation analysis of gene level sum indel frequency in GBM mice induced with AAV-mTSG library

Table S19. Brain tumorigenesis statistics of mice injected with AAV sgRNA minipools targeting various genes or combinations, and controls

Table S20. Metadata for RNA-seq with mouse autochthonous GBM cells with various genotypes and treatments

Table S21. RNA-Seq estimated count table

Table S22. Differential expression analysis of *Rb1* vs. *Nf1* cells

Table S23. Differential expression analysis of *Nf1;Mll3* vs. *Nf1* cells

Table S24. Differential expression analysis of *Rb1;Zc3h13* vs. *Rb1* cells

Table S25. Differential expression analysis of *Rb1* cells, TMZ vs. DMSO

Table S26. Differential expression analysis of *Rb1;Pten* cells, TMZ vs. DMSO

Table S27. Differential expression analysis of *Rb1;Zc3h13* cells, TMZ vs. DMSO

Table S28. Venn diagram of genes downregulated after TMZ treatment in *Rb1*, *Rb1;Pten*, and *Rb1;Zc3h13* cells.

Table S29. Venn diagram of genes upregulated after TMZ treatment in *Rb1*, *Rb1;Pten*, and *Rb1;Zc3h13* cells.

Table S30. Metadata for exome sequencing of select PBS, AAV-vector, and AAV-mTSG brain samples

Table S31. Merged and filtered exome variant calls in mTSG brain samples

Table S32. Survival statistics of mTSG, vector or PBS treated mice in the lentiCRISPR direct *in vivo* GBM screen

Table S33. Primers used for T7E1 assays to assess sgRNA cutting

